# Identification of a novel GREMLIN1 uptake pathway in epithelial cells that requires BMP binding

**DOI:** 10.1101/2025.01.09.632157

**Authors:** Zhichun Gao, Yuhan Gao, Louise R. Dutton, Grace Todd, Gregory R. Gipson, Connor Browne, Emma M. Kerr, Carole Daly, Bianca Plouffe, Philip D. Dunne, Derek P. Brazil

**Affiliations:** Wellcome-Wolfson Institute for Experimental Medicine, School of Medicine, Dentistry, and Biomedical Sciences, Queen’s University Belfast, 97 Lisburn Road, Belfast BT9 7BL, Northern Ireland, UK; Institute of Cancer Sciences, Wolfson Wohl Cancer Research Centre, University of Glasgow, Bearsden, Glasgow G61 1BD, Scotland, UK; Cardiovascular Research Institute, Massachusetts General Hospital, Harvard Medical School, 149 13^th^ Street, CNY149-4100, Boston MA 02129, USA; Patrick G. Johnston Center for Cancer Research, School of Medicine, Dentistry, and Biomedical Sciences, Queen’s University Belfast, 97 Lisburn Road, Belfast BT9 7BL, Northern Ireland, UK

## Abstract

Gremlin1 is a member of a cysteine-knot containing family of secreted antagonists of bone morphogenetic protein signaling. GREM1 binding to BMP targets prevents their engagement with cognate BMP receptors, attenuating BMP-dependent gene expression. Some evidence suggests that GREM1 can directly bind to receptor tyrosine kinases on the plasma membrane, further complicating our understanding of GREM1 biology. To attempt to clarify the modalities of GREM1 signaling, we show that GREM1 protein is produced and secreted by intestinal fibroblasts and endocytosed by neighbouring epithelial cells. GREM1 uptake is a slow process and occurs by both clathrin- and caveolin-mediated endocytosis. Cell membrane heparin sulfate proteoglycans are required for GREM1 binding and uptake, and once internalised, GREM1 appears to localise to the early endosomes. Addition of BMP2 enhanced GREM1 uptake into cells. Remarkably, generation of a BMP-resistant GREM1 mutant abolished GREM1 uptake both in the presence and absence of BMP2. These data suggest that GREM1 binding and uptake into cells requires BMP binding, a process that may contribute to the antagonism of BMP signaling by GREM1.

**Summary:** In this article, we demonstrate differential GREM1 mRNA versus protein expression in mouse intestine. We also identify a novel GREM1 endocytosis pathway whereby mammalian cells take up GREM1 protein in what appears to be a BMP-dependent mechanism.

## Introduction

Gremlin1 (GREM1) is a secreted glycoprotein antagonist of bone morphogenetic proteins. GREM1 binds to BMP targets such as BMP2 and BMP4 to sequester them in the extracellular matrix, preventing BMP-mediated receptor binding and activation (Brazil et al., 2015). Exquisite temporospatial control of GREM1 and BMP expression in required for normal mammalian limb and kidney development. GREM1 is the BMP antagonist required for maintenance of the apical ectodermal ridge (AER) via regulation of sonic hedgehog (Shh) and fibroblast growth factor (Fgf) signaling during limb development (Khoka et al., 2003), (Zuniga et al.,1999), (Panman et al., 2006) Mice with homozygous deletions in the *Grem1* gene display defective fore and hindlimb formation, as well as renal agenesis, leading to death shortly after birth (Khoka et al., 2003), (Zuniga et al.,1999). These defects are thought to be due to inappropriately amplified BMP4 signaling, as deletion of a single allele of Bmp4 can rescue ureteric bud outgrowth and kidney morphogenesis in *Grem1*-/- mice (Michos et al., 2007). GREM1 expression in adult tissues has been identified mainly in stromal fibroblasts. For example, GREM1 expression in a distinct population of fibroblasts was shown to play a key role in intestinal regeneration after injury (Koppens et al., 2021). GREM1 expression has also been described in a range of stem cells including intestinal mesenchymal stem cells (Hong et al., 2018), osteochondroreticular (OCR) cells (Worthley et al., 2015) and glioma cancer stem cells (Yan et al., 2014). GREM1 plays a key physiological role in essential morphogen gradients required for normal tissue development and homeostasis.

Along with its critical role in development, GREM1 upregulation has been reported in a range of human pathologies. Elevated GREM1 mRNA in samples from patients with idiopathic pulmonary fibrosis (Koli et al., 2006), (Myllärniemi et al., 2007), diabetic nephropathy (Dolan et al., 2005) chronic pancreatitis (Staloch et al., 2015) and osteoarthritis (Chang et al., 2019). GREM1 overexpression has also been detailed in a wide range of human cancers, including colorectal (Tomlinson et al., 2011), (Davis et al., 2015), mesothelioma (Wang et al., 2011), gastric (Yamasaki et al., 2018), (Sun et al., 2020), breast (Sung et al., 2020), (Kim et al., 2020) and glioma (Guan et al., 2017), (Yan et al., 2014). A rare inherited condition in Ashkenazi Jewish families called hereditary mixed polyposis syndrome (HMPS) is caused by a 40 kb chromosomal duplication event on chromosome 15q13.3 upstream of the GREM1 gene that leads to 2500-fold upregulation of GREM1 mRNA production in the intestine of these patients (Jaeger et al., 2012). A mouse model of GREM1 overexpression in intestinal epithelial cells recapitulated this proliferative phenotype, suggesting that high levels of GREM1 alone can drive intestinal cell growth, intestinal polyp and tumor formation (Davis et al., 2015).

Most data in the literature suggest that GREM1 is a “bad actor” in human cancer, with high levels of GREM1 associated with worse patient outcomes (e.g. (Davis et al., 2015), (Dutton et al., 2019). However, a small number of papers argue the opposite, and suggest that high levels of GREM1 is a prognostic marker of less aggressive tumor phenotypes and improved patient outcomes in both pancreatic adenocarcinoma (Lan et al., 2022) and colorectal cancer (Jang et al.,2016), (reviewed in (Gao et al., 2023)).

In addition to its canonical role as a secreted ligand-trap antagonist for BMP2 and other BMPS, some reports have suggested a non-canonical, direct signaling capacity for GREM1. GREM1 was reported to bind and activate the vascular endothelial growth factor receptor-2 (VEGFR2), initiating angiogenesis (Mitola et al., 2010), (Chiodelli et al., 2011). However, other groups including our own reported conflicting data on the ability of GREM1 to activate VEGFR2 (Dutton et al., 2019), (Rowan et al., 2018). A recent paper in Nature reported that GREM1 could activate the fibroblast growth factor receptor 1 (FGFR1) to activate MEK/ERK signaling and drive castration-resistant prostate cancer growth (Cheng et al., 2022). Together with other reports that GREM1 can activate epidermal growth factor (EGF) receptor in breast cancer cells (Park et al., 2020), (Sung et al., 2020) and bind to slit guidance ligand-2 (SLIT2) in neurons, a somewhat muddied picture exists of GREM1 biology and signaling in health and disease. To address some of the gaps in our understanding of GREM1 biology, we have interrogated the secretion, membrane binding and uptake of GREM1 into mammalian cells. We report that GREM1 is secreted, binds to, and is endocytosed in colorectal cancer cells, a process that requires BMP binding.

## Methods

### Cell Culture

HEK293T were cultured in Dulbecco’s Modified Eagle Medium (DMEM) containing pyruvate, 1 g/L glucose (Gibco, UK, Cat. No: 31885023), supplemented with 10 % FBS (Gibco^TM^, UK, Cat. No: 10082147) and 1 mM HEPES (SIGMA-ALDRICH, Cat. No: H0887). HEK293Tcells were maintained in T75 cm² flask at 37 °C, 5 % CO_2_ and plated on 6-well plates or 10 cm^2^dishes for treatments. HCT116 human colon cancer cells were cultured in McCoy’s 5a Medium Modified with L-Glutamine (Gibco^TM^, UK, Cat. No: 26600023), supplemented with 10 % FBS (Gibco^TM^, UK, Cat. No: 10082147) and 1 mM sodium pyruvate (Gibco^TM^, UK, Cat. No: 11360070). HCT116 cells were maintained in T75 cm² flask at 37 °C, 5 % CO_2_. HeLa cells were cultured in DMEM containing low glucose, pyruvate, and L-glutamine (Gibco^TM^, UK, Cat. No: 31885023), supplemented with 1x non-essential amino acids (Gibco^TM^, UK, Cat. No: 11140035) and 10 % FBS (Gibco^TM^, UK, Cat. No: 10082147). HeLa cells were grown in T75 cm² flask at 37 °C, 5 % CO_2_. C2C12-BRE (BMP response element) cells were a generous gift from Professor Gareth Inman (Beatson Institute for Cancer Research, University of Glasgow). Cells were maintained in DMEM (Gibco^TM^, UK, Cat. No: 31885023) supplemented with 10 % FBS (Gibco^TM^, UK, Cat. No: 10082147) in T175 cm^2^ flasks at 37 °C in a humidified 5 % CO_2_ incubator. All cell lines were screened every month and confirmed as negative for *Mycoplasma* using a commercially available PCR test kit.

### HEK293 Transfection

HEK293T cells were transfected when they reached 60 % confluence in 10 cm^2^ dishes well plates. Each dish was transfected with 600 μL of Opti-MEM^TM^ (Gibco^TM^, UK, Cat. No: 31985047) and 18 μL Lipofectamine 2000 Invitrogen, UK, Cat. No: 11668019) transfection reagent per dish. The appropriate volume of Lipofectamine 2000 and Opti-MEM^TM^ MasterMix were mixed followed by 5 min incubation at RT. Then 6 µg of plasmid per dish were added along with 600 µL of Opti-MEM in MasterMix followed by 20 min incubation at RT. An additional 12 mL of DMEM supplemented with 10% FBS were added to the final mixture media before adding to cells. Medium was changed after 5 h incubation at 37 °C and cells were harvested after 24 h or 48 h post-transfection.

### Protein Extraction and Western Blotting

Total protein was extracted using RIPA buffer (50 mM Tris-HCl, pH 7.4, 0.5 % (v/v) sodium deoxycholate, 150 mM sodium chloride (NaCl, Melford, UK, Cat. No: S23020), 1 % (v/v) Triton™ X-100 Surfact-Amps™ Detergent Solution (Triton X ThermoFisher Scientific, Cat. No: 85111) and 1 mM EDTA (SIGMA-ALDRICH, Cat. No: E5134). Prior to adding to cells, RIPA buffer was supplemented with 250 μM sodium orthovanadate (Na3VO4, SIGMA-ALDRICH, Cat. No: 450243), 40 mM β-glycerolphosphate (SIGMA-ALDRICH, Cat. No: 50020), 1 mM sodium fluoride (NaF, SIGMA-ALDRICH, Cat. No: 450022), 2 μM microsystin-LR (Enzo Life Sciences, Cat. No: ALX-350-012), 1 mM phenylmethanesulfonylfluoride (PMSF, SIGMA-ALDRICH, Cat. No: 329-98-6) and 1 x protease inhibitor cocktail (SIGMA-ALDRICH, Cat. No: P8340). Cell lysates were diluted with 2 × Laemmli buffer containing β-Mercaptoethanol before boiling at 95 °C for 5 min. Cell lysates were loaded into 10 % or 15 % (v/v) SDS-PAGE gels. Proteins were transferred to PVDF membranes which were blocked with 3 % BSA/TBST for 1h. Membranes were incubated in the various primary antibodies overnight at 4 °C (Anti-human Gremlin1 (R&D systems, Cat. No: AF956), Anti-mCherry antibody (Abcam, Cat. No: ab125096), Anti-β-actin (Cell Signalling, Cat. No: 3700S), Anti-PhosphoSMAD1(Ser463/465)/ SMAD5 (Ser463/465)/ SMAD8 (Ser465/467) (Cell Signalling, Cat. No: 13820S), Anti-SMAD1 (Cell Signalling, Cat. No: 6944S). Membranes were then washed with 3 times with 1 X TBST and incubated with respective secondary antibody (1:10,000) for 1 h at RT. After 3 washes in 1 X TBST, membranes were exposed to enhanced chemiluminescence (ECL) reagents (Merck, Cat. No: WBKLS0050), and bands were visualised using the Genesys (G: BOX) imaging system.

### Immunofluorescence

Cells were seeded on either plastic plates or glass coverslips. Experimental treatments were initiated when cells were at 30 %-40 % confluency. Once fixed using 4 % (w/v) PFA (SIGMA-ALDRICH, Cat. No: 158127) or 10 % (w/v) formaldehyde (ThermoFisher Scientific, Cat. No: 28908), cells were permeabilized with 0.1 % Triton-X in PBS and blocked with PBS contain 0.5 % (v/v) Triton-X and 2 % (w/v) BSA for 2 h at 4 °C. Cells were incubated at 4 °C overnight in primary antibody (Anti-EEA1 1:100, Cell Signaling, Cat. No: 2411S), MitoTracker Red (Invitrogen, Cat. No: M7512), Anti-Calreticulin (Abcam, Cat No: ab92516), Anti-GM130 (1:500, BD Transduction Laboratories™, Cat. No: 610822), Anti-LAMP1 (1:50, Abcam Cat. No: ab25630), Anti-Rab 11a (1:100, ThermoFisher Scientific, Cat. No: 700184), made up in blocking buffer. After 3 x PBS washes, Alex-568 labelled secondary (rabbit anti-mouse) antibody was added at a 1:1000 dilution in blocking buffer, for 1 hour at RT. Cells were washed 3 x with PBS, and coverslips were then placed onto glass slides using VECTASHIELD^®^ Mounting Medium with DAPI (Cat. No: H1200) and left to dry overnight RT protected from light until imaging.

### Immunohistochemistry

FFPE sections (5 μm) were deparaffinized in xylene and rehydrated through a graded ethanol series. Endogenous peroxidase activity was quenched by incubating the sections in 3 % (v/v) hydrogen peroxide in PBS for 20 min. Antigen retrieval was conducted by heating the sections in 10 mM sodium citrate buffer (pH 6.0), followed by cooling at RT. Tissue sections were then permeabilized and blocked with 1 % (v/v) normal rabbit serum (Vectastain Elite ABC kit, Vector Laboratories, Cat. No: PK6105) in 2.5 % BSA – 0.3 % Triton-X PBS. To further minimize non-specific binding, an Avidin/Biotin Blocking Kit (Vector Laboratories, Cat. No: SP2001) was applied as per the manufacturer’s protocol. Sections were then incubated overnight at 4 °C with a primary anti-mouse GREM1 (1:200, R&D Systems, Cat. No: AF956) or an isotype control (goat normal IgG, 1:200, Santa Cruz, Cat. No: AB108C) in a humidified chamber. Following washing with 0.1 % PBS Tween, a secondary biotinylated rabbit anti-goat antibody (1:250, Vectastain Elite ABC kit, Vector Laboratories, Cat. No: PK6105) was added for 1 h at RT, followed by incubation with Streptavidin peroxidase (1:250, Vector Laboratories, Cat. No: SA5004) for 30 min. Visualization was performed using 3,3’-diaminobenzidine (DAB) substrate (DAB substrate kit, Abcam, Cat. No: ab64238), and the reaction was terminated in deionized water. Sections were counterstained with haematoxylin for 15 sec, dehydrated through graded ethanol solutions, cleared in xylene, and mounted with DPX. Sections were scanned using Leica AT2 Aperio scanner and images captured using Leica Aperio ImageScope.

### GREM1-FITC Generation and Treatment

Fluorescein isothiocyanate (FITC)-labelled recombinant human (rh) GREM1 (R&D Systems, Cat. No: 555190-GR-050) was generated using a FITC Conjugation Kit (Abcam, Cat. No: ab102884). This resulted in a 183 μg/ml solution of GREM1-FITC. GREM-FITC was directly diluted into the cell culture medium with different concentrations used in specific experiments (0.5 μg/mL, 1 μg/mL). Cells were protected from light prior to imaging, and GREM1-FITC conjugates were stored at 4 °C.

### Conditioned Medium Treatments

After 24 h or 48 h of HEK293T cell transfection, conditioned medium (CM) was collected by washing the cells once with PBS, followed by the addition of serum-free medium for 5 h at 37 °C. The CM was centrifuged at 1,200 rpm (13684 *x g*) for 10 min to remove any floating cells or cellular debris. The CM was then either applied to fresh cells or snap-frozen in liquid nitrogen and stored at -80 °C for later use. Ten % FBS (Gibco^TM^, UK, Cat. No: 10082147) and recombinant human (rh) BMP2 (R&D system, Cat. No: 355-BM-050/CF) were employed to measure the effect on GREM1 internalization. The experiments were conducted in both HCT116 and Hela cells, and GREM1 was expressed as a secreted GREM1^WT^ or GREM1^Mut^-mCherry fusion protein. CM was supplemented with or without 10 % FBS, and then incubated with vehicle (4 mM HCl and 0.1 % BSA) or 200 ng/mL BMP2 overnight (16 h). Following incubation, cells were washed with PBS, fixed and stained with DAPI (Thermo Fisher Scentific, Cat. No: D1306) at a concentration of 1 µg/mL in PBS for 20min. . Imaging was performed using a DMi8 microscope at 20 x magnification.

### Fluorescence Quantification of GREM1-FITC and GREM1-mcherry

GREM1-FITC or GREM1-mCherry integrated cell fluorescence were measured using ImageJ. Images displaying only FITC or mCherry fluorescence were used as quantification controls. Prior to measurement, all images were blinded and the integrated intensity per cell was calculated with background subtraction. The sum of the integrated density within the selected regions was measured by ImageJ. The total number of cells in each image was counted, with partial cells at the image boundaries counted only on the top and left sides. The integrated fluorescence intensity per cell was then determined by dividing the total integrated density by the number of cells. After completing the quantification, the data were unblinded and subsequently analyzed and plotted using GraphPad Prism (Version 9.1.0).

### Endocytosis Inhibitor Treatments

HeLa or HCT 116 cells were cultured to 30 %-40 % confluency on coverslips in 24-well plates. Pitstop 2 (SIGMA-ALDRICH, Cat. No: SML1169) was dissolved in DMSO to achieve a stock concentration of 30 mM. Initially, cells were pre-incubated with 25 μM Pitstop 2 in serum free medium for 10 min, then washed once with PBS. After pre-incubation, 0.5 μg/mL GREM1- FITC were added to the cells in serum-free medium for another 6 h before fixing and imaging. Dyngo 4a (Abcam, Cat. No: ab120689) was dissolved in DMSO to a final stock concentration of 10 mM. Cells were initially pre-incubated in 30 μM Dyngo 4a in serum free medium for 30 min. After once PBS wash, cells were incubated with 0.5 μg/mL GREM1-FITC in serum free medium an additional 6 h before fixation and imaging. Heparinase lll (SIGMA-ALDRICH, Cat. No: H8891) was reconstituted in 20 mM Tris-HCl, pH 7.5, containing 0.1 mg/mL sterile BSA and 4 mM CaCl2 to a final stock concentration of 50 U/mL. Cells were initially pre-incubated in Heparinase lll in complete medium for 2 h. After pre-incubation, 0.5 μg/mL GREM1-FITC was added to the cells in serum-free medium overnight before fixing and imaging.

### PNGase treatment of GREM1

The removal of high mannose, hybrid, and complex oligosaccharides from GREM1-FITC was achieved using the PNGase F enzyme kit (Biolabs, Cat. No: P0704S). One µg of rhGREM1 or GREM1-FITC was combined with 2 µL of GlycoBuffer 2 (10 ×),100 U of PNGase F and H_2_O to make a 20 µL volume. The reaction mixture was gently mixed and incubated at 37 °C for 2 h. After stopping the reaction with 2 × Laemmli buffer and boiling at 95 °C, deglycosylation of GREM1-FITC was verified accelerated GREM1 protein migration on Western blot (WB). For cell treatments, a parallel reaction without PNGase F, referred to as control GREM1-FITC, was run. Deglycosylated GREM1-FITC internalisation experiments were conducted using HeLa cells that were seeded in a 12-well plate and grown to 30-40 % confluency. Cells were treated overnight with 1 µg/mL of either control or deglycosylated GREM1-FITC. Subsequently, cells were fixed, stained with DAPI, and imaged using a DMi8 microscope.

### Statistical analysis

Graphs were generated and statistical analysis was conducted using GraphPad Prism 5.0 software. Data were plotted according to the format to highlight interexperimental reproducibility and variability (Lord et al, 2020). The statistical significance of two-group comparisons was assessed using Student’s t-test, while comparisons involving three or more groups were analyzed using One-way ANOVA followed by Bonferroni post-hoc test. Each experimental group comprised a minimum of three samples and experiments were repeated at least n=3 times unless otherwise specified. *p* values were calculated and present as * *p* < 0.05; ** *p* < 0.01; *** *p* < 0.001.

## Supplemental Figures Methods

### Luciferase assay

C2C12-BREs were seeded at a density of 1.0 x 10^5^ per well, per 24 well plate in DMEM supplemented with 1 % FBS for 24 h. Compound dilutions were prepared in certain volume of normal media DMEM and CM. Following addition of 5 ng/mL rhBMP2 or PBS (vehicle) in triplicate, mixtures (300 μL/well) were added to C2C12-BREs at 37 °C and incubated for 5 h. Treatment medium was then removed, cells washed once with ice cold PBS and lysed in 100 μL cell culture lysis reagent. Lysates were centrifuged (13,300 × g, 5 min, 4 °C). 30 μL respective supernatants and an equal volume of Luciferase assay reagent (Promega, Cat. No: E4530) were transferred in duplicate to a 96-well micro plate. Luminescence was then measured by FLUOstar® Omega plate reader.

### Live cell image capture

HCT116 cells were seeded onto an 8-well Ibidi μ-Slide at a density of 4 x 10^5^ cells/mL (300 μL per well). The experiment was generated when the cells reached 30-40 % confluency. Cells were treated with 1 μg/mL GREM1-FITC and positioned in the imaging chamber. Live cell imaging was conducted using a Nikon 6D Live Cell Imaging Inverted Microscope. To replicate incubator conditions within the sample chamber, the temperature was set at 37 °C and CO_2_ at 5 %, adjusted 30 min prior to placing the cells in the chamber. After adjusting the position and focus for each well, images were captured every 30 min for 16 h.

## Results

*In situ* hybridisation (ISH) of wild-type and *GREM1*-/- mouse colon FFPE sections identified a staining pattern consistent with GREM1 mRNA localisation to the muscularis layer and submucosa (Fig. 1a). No *GREM1* mRNA was detected in the colonic crypts, epithelial or other cells (Fig. 1a). The specificity of the ISH probes was confirmed by the absence of any positive signal in FFPE sections from *GREM1*-/- mice (Fig. 1b). In contrast, GREM1 protein was more widely detected in mouse colon, with staining detected in epithelial cells at the base of the crypts, as well as in the muscularis and submucosa layers (Fig. 1c). Important concerns around the specificity of the anti-GREM1 antibody used for immunohistochemistry staining were allayed by the absence of any GREM1 protein signal in colon tissue from *GREM1*-/- mice (Fig. 1d). These results are consistent with previous data from our group characterising GREM1 mRNA/protein localisation in mouse intestine (Dutton et al., 2019). To extend these findings to a colorectal cancer model, colon sections from *Apc^fl/fl^;Kras^Lsl-G12D^;Tp53^fl/fl^;villin- CreERT2* (Shibata et al.,1997), (Jackson et al., 2001), (Marino et al., 2000) mice were probed for GREM1 protein expression. GREM1 protein expression was detected in the muscularis and submucosa layers of the colon, as well as intestinal crypts at the base of the tissue (Fig. 1g). Interestingly, expression of GREM1 protein was also detected in AKP colonic tumours, with strong staining at the basolateral surface of colonic epithelial cells (Fig. 1h). Little or no signal was obtained using isotype control antibodies (Fig. 1e, f). These data suggest that low levels of GREM1 protein are detected in normal mouse intestine, with higher levels evident in AKP intestine and tumour tissue.

**Figure 1.** Differential pattern of GREM1 mRNA and protein expression in healthy and colorectal cancer intestine. FFPE colon sections (5 μm) from *wild-type* or *GREM1*-/- mice were analyzed for GREM1 mRNA and protein expression. *In situ* hybridisation (a, b) and immunohistochemistry (c. d) were performed as described in Methods. FFPE colon sections (5 μm) from *Apc^fl/fl^;Kras^Lsl-G12D^;Tp53^fl/fl^;villin-CreERT2* (AKP) mice were stained for GREM1 protein (e-h) as described in Methods. Images from both mouse intestine (e, g) and tumor (f, h) were captured after staining with goat IgG isotype control (e, f) or anti-GREM1 antibody (g, h). Positive GREM1 protein staining was visualised using DAB (brown), and sections were counterstained with haemotoxylin (blue) and imaged using PathXL and Aperio ImageScope. Scale bars 100 μm (a-d), 200 μm (e, f) or 300 μm (g, h). Images are representative of staining from n=4 mice.

Based on these results, we hypothesised that fibroblasts or other cells in the muscularis layer of the intestine express *GREM1* mRNA that is translated to GREM1 protein which is secreted and taken up by proximal epithelial cells. To interrogate this hypothesis, GREM1-FITC protein was generated and added to HeLa cells. GREM1-FITC was indistinguishable from non-labelled GREM1 in terms of their ability to inhibit BMP2-mediated SMAD1/5/8 phosphorylation (Suppl. Fig. 1A). At 60 min, little or no uptake of GREM1-FITC, or control FITC-BSA was evident (Fig. 2 a-f). In contrast, overnight incubation led to clear uptake of GREM1-FITC, but not BSA-FITC into HeLa cells (Fig. 2 g-l). Similar data were obtained using HEK293 cells (data not shown). High resolution confocal images demonstrated a distinct punctate, perinuclear staining pattern for GREM1-FITC, with staining also detected in cellular extensions and contact points between cells (Fig. 2 m, n). Having established the GREM1-FITC uptake assay in HeLa cells, we then measured the time-course of GREM1-FITC uptake into HCT116 human colonic epithelial cells (Fig. 3). Binding of GREM1-FITC to HCT116 cell membranes became evident between 5-15 min (Fig. 3), with a ring-like membrane-bound pattern evident at 60 min (Fig. 3). Internalisation of GREM1-FITC began at 3-6 h, with most of the GREM1-FITC detected inside the cells between 16-24 h (Fig. 3). Live-cell imaging of GREM1-FITC uptake into HCT116 cells supported these conclusions (Suppl. Fig. 2 and video). These data suggest that GREM1 internalisation is a slow process that may involve GREM1 binding to membrane proteins or receptors prior to internalisation.

**Figure 2.** GREM1 is internalized by HeLa cells. HeLa cells were treated with PBS, 0.5 μg/mL BSA-FITC or GREM1-FITC in complete medium for 60 min or overnight. After incubation, cells were washed with PBS to remove any free FITC conjugates and fixed with 4 % PFA and DAPI stained before imaging using DMi8 as described in Materials and Methods. Panels show FITC fluorescence (green) and overlay with phase contrast and DAPI staining (nucleus, blue) (a-l). Higer magnification images using Leica SP5 Confocal laser scanning microscope is shown in m,n. GREM1-FITC located in cell extensions is indicated with arrow. Data are representative of n=5 independent experiments.

**Figure 3.** GREM1 uptake is slow and involves accumulation at the plasma membrane. HCT116 cells were treated with 1 μg/mL GREM1-FITC for the indicated times before washing once with PBS, fixation with 4 % PFA and DAPI staining as described in Methods. FITC and DAPI staining were then visualized on a Leica SP5 Confocal laser scanning microscope. Scale bar, 25 μm and is shown on right hand side.

Several reports have identified that heparin sulphate proteoglycans (HSPGs) are involved in GREM1 signalling and biology (Chiodelli et al., 2011), (Pegge et al., 2020), (Tatsinkam et al., 2015). Removal of heparin moieties by cell treatment with Heparinase III significantly reduced GREM1-FITC uptake, suggesting that binding or interaction with HSPG-containing membrane proteins is required for GREM1 uptake (Fig. 4A, B). GREM1 is also a glycoprotein, with the role of glycosylation suggested to be cell association and binding of GREM1 to the extracellular matrix (Topol et al., 2000). PNGase deglycosylase treatment of GREM1-FITC led to a downward shift in mobility on SDS-PAGE, confirming deglycosylation (Fig. 4C). This deglycosylated form of GREM1-FITC was still able to inhibit BMP2-mediated SMAD1/5/8 phosphorylation (data not shown). Deglycosylated GREM1-FITC was taken up in HeLa cells to the same extent as glycosylated GREM1-FITC (Fig. 4D, E), suggesting a non-critical role for GREM1 glycosylation for it’s uptake. These data suggest that while heparin-containing proteins such as HSPGs are required for GREM1 internalisation, glycosylation of GREM1 does not appear to be a critical mediator of it’s binding and uptake into cells.

**Figure 4.** Heparin sulfate proteoglycans, but not glycosylation is required for GREM1 uptake into cells. A. HeLa cells were preincubated with vehicle or Heparinase III (0.5 U/mL) for 2 h, followed by addition of 0.5 μg/mL GREM1-FITC overnight. C. GREM1-FITC (500 ng) was treated with vehicle or 50 U PNGase F for 2 h, with deglycosylation confirmed by Western which were then added to HeLa cells overnight (D, E). Cells were fixed in 4 % PFA and cell nuclei were stained with DAPI as described in Methods. FITC and DAPI images were captured using a DMi8 Leica Microscope at 20 x magnification. B, E. Integrated fluorescence per cell was calculated using ImageJ for 3 images per technical replicate, with n=3 independent experiments plotted. Large circles, squares, and triangles represent the mean fluorescence intensity from each experiment, and smaller symbols represent the fluorescence intensity of each technical replicate within each individual experiment. Statistical significance was determined using a two-tailed Student’s t-test (NS, non-significant; *, p < 0.05).

To further characterize the mechanisms of GREM1 internalisation, HeLa cells were treated with PitStop2, an inhibitor of clathrin-mediated endocytosis (Robertson et al., 2014). After 6 h incubation with GREM1-FITC, HeLa cells displayed a predominantly intracellular staining pattern of GREM1-FITC (Fig. 5Ai, ii). In contrast, cells treated with Pitstop2 demonstrated less intense staining of GREM1-FITC, with lower intracellular levels evident (Fig. 5Aiii, iv). Consistently, when HCT116 cells were used, more intense GREM1-FITC staining was detected at the cell-cell junctions (i.e. extracellular) in Pitstop2-treated cells, compared to a more intracellular pattern of staining in control cells (Fig. 5Bi-iv). Quantification demonstrated an approximate 40 % decrease in GREM1-FITC uptake in HeLa cells (Fig. 5C). These data suggest that at least some GREM1 uptake into cells is occurring via clathrin-mediated endocytosis. Treatment of HeLa cells with Dyngo4a, a dynamin inhibitor of caveolin-mediated endocytosis (Macia et al., 2006) reduced GREM1-FITC uptake by ∼80 % (Fig. 5D, F). A similar result was obtained when HCT116 colonic epithelial cells were utilized, with most of the GREM1-FITC staining detected at the cell-cell junctions in Dyngo4a-treated cells (Fig. 5Eiii, iv), with little or no intracellular GREM1-FITC evident compared to control cells (Fig. 5Ei, ii). Together, these data suggest that both clathrin and caveolin-mediated endocytosis are involved in GREM1 uptake into cells, with a more prominent role for caveolin-mediated pathways.

**Figure 5.** GREM1 internalization occurs via clathrin and caveolin-mediated endocytosis. HeLa (A) or HCT116 (B) cells were pretreated with DMSO or 20 μM Pitstop® 2 for 10 min and further incubated with 0.5 μg/mLGREM1-FITC for 6h. HeLa (D) or HCT116 (E) cells were also pretreated with vehicle (DMSO) or 30 μM Dyngo 4a for 30 min then incubated with 0.5 μg/mL GREM1-FITC for 6h. Cells were fixed in 4 % (w/v) PFA and cell nuclei were stained with DAPI as described in Methods. FITC and DAPI were imaged using a DMi8 Leica Microscope at 20 x magnification (A,D) or an SP5 confocal microscope at 100 x magnification with crop action (B,E) . Scale bar, 50 μm and is shown on left hand side. Integrated Fluorescence per cell treated with Pitstop^®^ (C) or Dyngo 4a (F) was calculated using Image J for 3 images per technical replicate with n=3 independent experiments plotted. Large circle, square and triangle represent the mean fluorescence intensity within each experiment (biological replicate) and small symbols represent the fluorescence intensity of each image within each individual experiment (technical replicates). Statistical significance was calculated using a two-tailed Student’s t-test, **, p<0.01.****, p<0.0001.

Confocal microscopy identified a distinct pattern of staining for GREM1-FITC inside cells, with a punctate, perinuclear pattern evident (Fig. 2, 3), and evidence of GREM1-FITC in filopodia and other cellular extensions (Fig. 2m, n). No significant localisation of GREM1-FITC to either mitochondria (stained with MitoTracker) or ER (stained with calreticulin) were detected (Suppl. Fig. 3), with very slight staining evident in the Golgi apparatus using GM130 and giantin as Golgi markers (Suppl. Fig. 3C and data not shown). The punctate staining pattern was suggestive of lysosomal or endosomal localisation. A small amount of GREM1-FITC staining was detected in lysosomes (identified using anti-LAMP1 or cresyl violet staining, Suppl. Fig. 4 and data not shown). Staining of cells with EEA1, a marker of early endosomes, demonstrated some overlap with GREM1-FITC staining in HeLa cells (Fig. 6A). These data suggest that internalised GREM1 partially localises to the early endosomal compartment. The hypothesis that internalised GREM1-FITC could be recycled and resecreted from cells was then tested. Treatment of fresh HeLa cells with CM captured from GREM1-FITC treated cells generated green cells (Fig. 6B), suggesting that endocytosed GREM1-FITC could be resecreted and taken up into new cells after resecretion. However, no significant overlap in staining of GREM1-FITC was detected when Rab11 was used a marker of recycling endosomes (Suppl. Fig. 4B), suggesting that resecretion via this pathway may not be the primary mechanism for intracellular GREM1 recycling.

**Figure 6.** GREM1 can be resecreted from cells after endocytosis and localizes to the early endosomes. A. HeLa cells were seeded on glass coverslips in 24-well plates. After adherence, cells were treated with 0.5 μg/mL GREM1-FITC (green) overnight in complete medium. Cells were then washed with PBS and fixed with 4 % PFA (w/v) before staining with anti-EEA1 (red) and DAPI (blue) to visualize the early endosomes and nuclei respectively. Slides were then images at 100 x magnification on a Leica SP5 confocal microscope. Scale bars represent 25 μm or 10 μm in magnified images. Images are representative of n=3 independent experiments carried out in duplicate. B. Conditioned medium (CM) was collected from HeLa cells treated with either FITC-BSA (1 μg/mL) or GREM1-FITC (1 μg/mL) overnight. Fresh HeLa or HCT 116 cells were treated with either CM fraction overnight. Cells were fixed in 4 % (w/v) PFA and cell nuclei were stained with DAPI. FITC and DAPI fluorescence were imaged using a DMi8 Leica microscope at 20 x magnification. Data are representative of n=3 independent experiments performed in duplicate. Scale bars, 50 μm.

One potential limitation of our study may be the reliance on commercially available recombinant human GREM1 protein (from Biotechne/R&D Systems). To overcome this, we generated plasmids that expression a fusion protein of GREM1 with mCherry (GREM1- mCherry, Fig. 7). GREM1-mCherry was detected at ∼55 kDa in HEK293T cells and inhibited BMP2-stimulated SMAD1/5/8 phosphorylation, confirming functionality (Fig. 7A). Similar data were obtained when BMP4 was used to stimulate SMAD1/5/8 phosphorylation (Suppl. Fig. 1B). Serum-free conditioned medium collected from transfected HEK293T cells confirmed that GREM1-mCherry was secreted, similar to endogenous GREM1 (Fig. 7B). Incubation of HeLa and HCT116 cells with conditioned medium from mCherry control or GREM1-mCherry transfected cells demonstrated clear uptake of GREM1-mCherry in overnight-treated cells, with low levels of uptake in HCT116 cells, but not HeLa cells at 60 min (Fig. 7C, D). Importantly, the intracellular staining pattern observed with GREM1-mCherry was consistent with that observed with GREM1-FITC, suggesting biological consistency when both commercially sourced rhGREM1 and expressed, secreted, “homemade” GREM1-mCherry was employed (Fig. 2, 3, 7).

**Figure 7.** GREM1-mCherry inhibits BMP2 signaling and is internalized by mammalian cells. A. HEK293T cells transfected with either mCherry empty vector or hGREM1-mCherry were treated with vehicle or 5 ng/mL BMP2 at 37 °C for 1 h before proteins were extracted and analysed by Western blotting using antibodies reactive to GREM1, mCherry, pSMAD1/5/8 and total SMAD1. Membranes were reprobed for β-actin as loading control. B. Conditioned medium (CM) was collected from transfected cells and analysed by Western blotting using antibodies reactive to GREM1 and mCherry. Cell lysate from transfected HEK293T cells used as positive control (+). Data are representative of n=3 independent experiments. HeLa (C) or HCT116 (D) cells were exposed to CM from pCMV-mCherry or pCMV-GREM-mCherry- transfected cells for 1 h or overnight (16 h). Cells were fixed with 4 % PFA, stained with DAPI, and imaged using the Leica Sp5 confocal microscope at 100 x magnification. Scale bars are indicated in the images. Images are representative of 6 images per triplicate technical replicate from n=3 independent experiments.

Previous reports have suggested that BMP2 could be transferred between neighbouring cells via vesicular transport, and that BMP antagonists such as Noggin increased BMP2 transfer between cells (Alborzinia et al., 2013), (Alborzinia et al., 2016). Addition of BMP2 to CM-containing GREM1-mCherry increased GREM1-mCherry uptake into HCT116 cells ∼5-fold (Fig. 8A, B). These data suggest that the quaternary complexes of GREM1 and BMP2, which are predicted to form fibril-like open-ended oligomers with interacting ⍺-helixes of GREM1 and BMP2 (Nolan et al., 2016), (Kišonaitė et al., 2016) may be more efficiently endocytosed by cells compared to non-BMP bound GREM1.

**Figure 8.** GREM1 endocytosis is increased by BMP2. A. CM from HEK293T cells transfected with GREM1-mCherry was supplemented with 10 % FBS before adding to HCT116 cells treated with vehicle (4 mM HCl and 0.1 % BSA) or 200 ng/mL BMP2 overnight (16 h). Cells were fixed and stained with DAPI before imaging using a Leica DMi8 microscope at 20 x magnification. B. Integrated fluorescence intensity per cell was quantified using ImageJ after blinding the images. Data are presented as mean ± SEM. Large symbols indicate the mean fluorescence intensity from n=3 independent experiments. Small symbols are representative of the average fluorescence of three images taken per technical replicate. Statistical analysis was carried out using a two-tailed Student’s t-test. ***, p<0.001

Previous research by Nolan et al identified that 3 specific amino acids (F125/I127/F138) were required for robust binding of GREM2 (another related BMP antagonist of the DAN family with 60 % sequence identity to GREM1) to BMP/GDF targets. Sequence alignment of GREM1 and GREM2 demonstrated that these amino acids are conserved in GREM1 and cluster together in β-sheet strands 2 and 3 (Fig. 9A). These 3 amino acids were mutated to Ala to generate a F125A/I127A/F138A GREM1 mutant (GREM1^Mut^) that were expressed, along with wild-type GREM1, as mCherry fusion proteins (GREM1^WT^-mCherry, GREM1^Mut^-mCherry, Fig. 9). A model of GREM1 binding to BMP2 generated using AlphaFold3 displays the effect of these 3 amino acid mutations on the putative binding surfaces between GREM1 and BMP2, suggesting a disruption of stabilizing hydrophobic interactions (Fig. 9B). Both GREM1^WT^- mCherry and GREM1^Mut^-mCherry expressed in HEK293 cells, with lower levels of GREM1^Mut^- mCherry detected in cell lysates, but higher levels detected in the CM, suggesting more efficient secretion of this GREM1 mutant (Fig. 9C). Addition of BMP2 demonstrated a clear increase in SMAD1/5/8 phosphorylation in control cells that was completely inhibited in GREM1^WT^-mCherry transfected cells (Fig 9C). In contrast, cells transfected with GREM1^Mut^- mCherry demonstrated little or no inhibition of BMP2-induced Smad1/5/8 phosphorylation (Fig. 9C). A similar reduction in the ability of CM containing GREM1^WT^ v GREM1^Mut^-mCherry to inhibit BMP2 was observed when C2C12 cells transfected with a BMP response element (BRE) luciferase reporter (Herrera et al., 2009) were utilised (Suppl. Fig. 1C).

**Figure 9.** A mutant GREM1 that cannot bind BMP2 displays reduced internalization. A. Sequence alignment of human GREM1 and GREM2 (*black text*) with identity and homology indicated (*blue text*). Key GREM1 amino acids identified for BMP binding (F125, I127 and F138) indicated in red. These amino acids cluster in the central two β-strands, with F125 and I127 at the end of strand 2 and F138A close to the start of strand 3 (Marco and Tom refs). All 3 of these amino acids were mutated to Alanine in the GREM1 mutant. B. Alpha Fold 3 predicted structures of GREM1^WT^ and GREM1^Mut^ bound to BMP2 with the positions of the 3 amino acids F125 (Phe101), I127 (Ile103) and F138 (Phe114) in the β-strands indicated. The numbering of the amino acids differs in the AlphaFold 3 structure as the mature GREM1 protein lacking the secretion signal (amino acids 1-24) was used to generate the predicted structure. C. HEK293T cells were transfected with plasmids expressing mCherry, GREM1^WT^- mCherry and GREM1^MUT^-mCherry. After 24 h transfection, cells were treated with vehicle (4 mM HCl and 0.1 % BSA) or 5 ng/mL BMP2 for 1 h before adding 1 ml serum-free media for 4 h to harvest conditioned medium (CM). Protein extracts were analyzed by Western blot, using antibodies reactive to pSMAD1/5/8, total SMAD1 and hGREM1. β-actin was used as a loading control. D. CM collected from HEK293T cells transfected with plasmids expressing mCherry, GREM1^WT^-mCherry or GREM1^MUT^-mCherry was added to HeLa cells overnight. Cells were fixed with 4 % PFA and DAPI stained before imaging (n=3 independent experiments). E. CM was collected and analyzed by SDS-PAGE. GREM1^WT^-mCherry or GREM1^MUT^-mCherry were detected by anti-hGREM1 antibody before addition of CM to cells (Input) or after overnight incubation (post-incubation). Images are representative of n=3 independent experiments carried out in triplicate. F. Image J was used to calculated GREM1 band intensities, and the ratio of GREM1 in post-incubation/input samples was calculated to estimate the percentage of free GREM1 remaining in the CM. Symbols represent means of triplicate values from n=3 independent experiments with error bars indicating SEM. Statistical significance was calculated using a two-tailed Student’s t-test *, p<0.05.

These data suggest that GREM1^Mut^-mCherry is resistant to BMP2 binding and may therefore be used as a tool to determine the requirement of BMP binding for GREM1 action. The uptake of GREM1^WT^-mCherry versus GREM1^Mut^-mCherry into cells was then compared. Addition of CM containing GREM1^WT^-mCherry to HeLa cells demonstrated clear uptake after overnight incubation (Fig. 9D). Surprisingly, uptake of GREM1^Mut^-mCherry was much lower compared to GREM1^WT^-mCherry (Fig. 9D). The lower intensity of mCherry fluorescence seen with GREM1^Mut^-mCherry was not caused by protein degradation, as GREM1^Mut^-mCherry protein was detected in the CM after removal from cells post-incubation at a similar level to the pre-incubation CM (Fig. 9E, F). In contrast, lower levels of GREM1^WT^-mCherry were detected in post- versus pre-incubation CM, likely due to partial uptake into cells (Fig. 9E, F). Addition of BMP2 significantly increased GREM1^WT^-mCherry uptake into HCT116 cells, but had no effect on GREM1^Mut^-mCherry uptake, consistent with an inability of this mutant form of GREM1 to bind to BMP2 (Fig. 10). These data support the conclusion that BMP binding is required for GREM1 uptake into mammalian cells.

**Figure 10.** BMP2 increases GREM1^WT^, but not GREM1^Mut^-mCherry uptake into HCT116 cells. A. Conditioned medium from HEK293 cells transfected with GREM1^WT^-mCherry or GREM1^Mut^-mCherry was supplemented with 10 % FBS and added to HCT116 cells in the presence of either vehicle (4 mM HCl and 0.1 % BSA) or 200 ng/mL BMP2 overnight for 16 h. Cells were fixed and stained with DAPI. Imaging was performed using Leica DMi8 microscope at 20 x magnification. Scale bars represent 50 μm. B. Integrated fluorescence intensity per cell was quantified and plotted by ImageJ and GraphPad Prism after blinding the images. Data are presented as mean ± SEM. Large symbols indicate the mean fluorescence intensity from n=3 independent experiments. Small symbols represent the average of duplicate wells, with three images taken per well. Statistical analysis was determined using one-way ANOVA followed by Bonferroni post-hoc test. (**, p<0.01; ***, p<0.001)

## Discussion

Our report identifies a novel mechanism involving the uptake of the secreted BMP antagonist GREM1 into cells after secretion, a process that is both enhanced by, and requires BMP binding. Canonical GREM1 signaling involves binding to BMP targets in the extracellular matrix leading to inhibition of BMP-mediated BMP receptor activation. Our report extends this model, suggesting that internalisation of the GREM1-BMP complexes may also contribute to attenuation of extracellular BMP signaling.

The pattern of *GREM1* mRNA and GREM1 protein expression in mouse intestine is unusual in that *GREM1* mRNA is detected in the muscularis and submucosa layers, with GREM1 protein also detected in these compartments, but also in adjacent epithelial and other cells in colonic crypts and intestinal villi (Fig. 1 and data not shown). These data suggest that the source of GREM1 protein in healthy intestine is fibroblasts which secrete GREM1 protein that is then taken up by proximal epithelial cells that do not contain *GREM1* mRNA. A consistent challenge with interpreting immunohistochemistry data for GREM1 is antibody specificity. In our hands, the most sensitive and specific anti-GREM1 antibody is from Biotechne/R&D Systems (AF956). The absence of GREM1 protein staining when intestinal sections from *GREM1*-/- wild-type mice were used increases the confidence in the positive GREM1 protein staining in these mice (Fig. 1d). Additional no primary antibody and normal goat IgG controls were included in the IHC staining of sections from AKP mice (Fig. 1e, f and data not shown). Again, low-to-absent signal for GREM1 protein in these controls suggests that positive DAB staining obtained for GREM1 accurately reflects the expression pattern of GREM1 protein in these tumor-bearing mice.

This disparity in mRNA versus protein expression detected from GREM1 has long been a challenge in biology. Genome-wide correlation between mRNA levels and protein expression are generally poor, with a figure of 40 % estimated in many reports (Koussounadis et al., 2015), (Vogel et al., 2012). This presents challenges when correlations between differentially expressed mRNA and their protein products are imputed from RNA sequencing data or quantitative PCR (Koussounadis et al., 2015). For example, a set of 17 immune and wound healing cytokines measured in 77 biopsies revealed a Spearman’s rank correlation of 0.67 when mRNA and protein abundance were measured in a model of canine wound healing (Prabahar et al., 2024). Liu and colleagues suggest that mRNA transcript levels alone are insufficient to predict protein levels in many cells and tissues, compromising our ability to predict genotype-phenotype relationships (Liu et al., 2016). Our data pose an important question: why is GREM1 gene transcription active in fibroblast/stromal cells, but not in epithelial cells? One obvious explanation is that the transcriptional landscape (including epigenetic marks of permissive chromatin such as H3K27Ac) is more favourable to GREM1 transcription in fibroblast versus epithelial cells. A second possibility is that transcription factors required to bind to cis-regulatory elements (CREs) on the GREM1 promoter are abundant in fibroblasts, but not present in epithelial cells. A third scenario is that transcriptional repressors may be present in epithelial cells that limit GREM1 expression compared to adjacent fibroblasts. To begin to unravel this conundrum, our group has identified cell-type specific transcription factors and their binding sites on GREM1 regulatory regions that regulate GREM1 mRNA production (Todd et al., in preparation). Further experiments will drive our understanding of the exact mechanisms of differential GREM1 expression in cells of the intestine.

A range of publications have detailed upregulated *GREM1* mRNA expression in a host of human cancers, including gastric (Sun et al., 2020), colorectal (Tomlinson et al., 2011), breast (Sung et al., 2020) and lung (Mulvihill et al., 2012). However, very few of these studies measured GREM1 protein expression in these cancers. Our data demonstrate that GREM1 protein expression is upregulated in the AKP transgenic mouse model of colorectal cancer, which express APC/KRAS/p53 mutations commonly found in human colorectal cancer (Shibata et al.,1997), (Jackson et al., 2001), (Marino et al., 2000). GREM1 protein was detected in the muscularis layer and epithelial cells of the colonic crypts, as well as in tumors that formed in these mice due to AKP transgene expression (Fig. 1). The triad of APC/KRAS/p53 mutations are commonly found in human colorectal adenocarcinoma. Our data suggest that upregulation of GREM1 protein is also a feature of tumors formed by these genetic alterations in mice. The next logical step would involve screening tissue microarrays from human colorectal cancers for GREM1 protein expression. However, currently available GREM1 antibodies show poor sensitivity and specificity for human GREM1, and existing data in the literature reporting GREM1 protein expression in human tissues should be interpreted with caution.

We describe for the first time the novel biological process of GREM1 protein uptake into epithelial cells Extensive characterization demonstrated that uptake of GREM1 was quite slow, with the majority of FITC-GREM1 outside the cell bound to the plasma membrane at the 3 h timepoint (Fig. 2, 3). Glycosylation of GREM1 was not necessary for internalisation, but removal of cellular HSPGs reduced GREM1 uptake by approximately 50 % (Fig. 4). Both clathrin and caveolin-mediated endocytotic pathways seem to be involved in GREM1 protein internalisation (Fig. 5). The pattern of GREM1 binding to the plasma membrane, together with the time-course and mechanism of endocytosis suggested that GREM1 may be binding to a membrane receptor prior to internalisation. Using a BRET-based approach monitoring activation of G-protein isoforms, GREM1 failed to activate any G-protein in either HEK293 or U251 glioblastoma cells, suggesting that GREM1 is not binding to a GPCR at the plasma membrane (data not shown).Reports in the literature have suggested that GREM1 can bind to receptor tyrosine kinases (RTKs) such as VEGFR2 (Mitola et al., 2010), EGFR (Park et al., 2020) and FGFR1 (Cheng et al., 2022), triggering downstream signal transduction. The reduction in GREM1 internalisation after Heparinase III treatment of cells suggests that some heparin sulfate-containing protein is involved in GREM1 binding at the plasma membrane. Data from the Rider laboratory identified the heparin-binding site of GREM1 (Tatsinkam et al., 2015) and showed that heparin sulfate binding was not essential for BMP antagonism (Tatsinkam et al., 2017). Given the high affinity of BMPs, particularly BMP2 and BMP4, for heparin sulfate, this may provide an alternative explanation for the decrease in GREM1 uptake after heparinase treatment. HSPG-bound BMPs are likely a localized reservoir for extracellular BMP, potentially serving to increase co-localization of GREM1 and BMP in the extracellular space, which may facilitate enhanced GREM1-BMP binding and subsequent internalization (Jiao et al., 2007), (Kanzaki et al., 2011), (Hettiaratchi et al., 2020), (Bramano et al., 2012). While RTKs such as FGFR1 signalling requires HSPGs (Ferguson et al., 2021), we should not exclude the possibility that other membrane HSPGs such as syndecans or glycosylphosphatidylinositol-anchored proteoglycans (glypicans) may be candidate receptors for GREM1 (Sarrazin et al., 2011).

The distribution of GREM1 when endocytosed appears as punctate, perinuclear staining in all cell lines tested (HEK293, HeLa, HCT116; Fig. 2). This pattern of staining was similar when both commercial recombinant GREM1 labelled with FITC or secreted GREM1-mCherry in conditioned medium was employed (Fig. 7). Our data suggest that endocytosed GREM1 does not localise exclusively to a specific subcellular compartment such as the Golgi or endoplasmic reticulum but seems to be distributed partially across the early endosomal and lysosomal compartments (Fig. 6A, Suppl. Fig. 4). An intriguing possibility is that GREM1 is associating in phase condensates, or membraneless intracellular compartments inside cells (Xu et al., 2024). We are currently testing this exciting hypothesis using chemical agents that disrupt phase condensate formation. GREM1 can be resecreted once taken up into mammalian cells (Fig. 6B) suggesting a dynamic environment exists in which GREM1 can be endocytosed and then resecreted for potential uptake into neighbouring epithelial cells in a paracrine-like manner. These data raise the question: what is the biological significance of GREM1 internalization by cells? Previous data demonstrated that prior to secretion, GREM1 can already bind precursor BMP4 and prevent mature BMP4 secretion, providing an intracellular mode of BMP antagonism (Sun et al., 2006). Given the somewhat slow rate of GREM1 endocytosis via both clathrin and caveolin-mediated pathways (Fig. 2) and the requirement for HSPGs for this process (Fig. 4A, B), the possibility exists that GREM1 cellular internalisation may occur as part of a complex with membrane receptors such as RTKs or other HSPG- containing proteins. Future experiments will be needed to identify whether GREM1 does indeed engage an RTK or other membrane protein, and whether this binding represents a previously unappreciated “non-canonical” signaling modality for GREM1, independent of BMP antagonism.

The complex of GREM1 with BMP2 was more efficiently endocytosed by HCT116 cells (Fig. 8). These data are supported by previous reports from the Wolfl group who demonstrated that BMP2 could be taken up into HeLa cells, and this uptake was increased by Noggin, but decreased by Chordin, both BMP antagonists (Alborzinia et al., 2013). Interestingly, this group demonstrated that lower molar excesses of GREM1 (2X and 5X) increased BMP2 uptake, but higher amounts (10X) had little or no effect (Alborzinia et al., 2013). These authors also demonstrate that Noggin can trap BMP2 inside the cell, and the authors propose a “transcytosis model of morphogen signalling” involving “….a trap-and-sink mechanism to modulate BMP signaling” (Alborzinia et al., 2016). Based on the enhancement of GREM1 uptake by BMP2, a similar model may exist for GREM1. However, it is not clear whether complexes of GREM1 and BMP2, once endocytosed, can be further dissociated inside the cell, thus allowing free GREM1 and BMP2 resecretion. Indeed, we propose the idea that the endocytosis of GREM1 as part of a complex with BMP targets may reduce the local concentration of BMP in the extracellular matrix, potentially leading to reduced extracellular BMP morphogen signaling.

The increase in GREM1 endocytosis in the presence of BMP2 would perhaps argue against the requirement for GREM1 to bind to a specific membrane receptor prior to internalization, assuming that BMP2 binding to GREM1 would reduce GREM1 binding to this putative receptor. The structure of the GREM1-BMP2 complex is predicted to be a collection of ‘fibril-like’ open-ended oligomers with interacting ⍺-helixes of BMP2 and GREM1 (Nolan et al., 2016), (Kišonaitė et al., 2016). One possibility could be that these large “daisy-chain”-like complexes more readily bind non-specifically to the plasma membrane, leading to enhanced endocytosis. However, we cannot exclude the possibility that an as-yet-unidentified heparin-containing membrane receptor may be involved in enhanced GREM1-BMP2 uptake. Interestingly, when these experiments were repeated in serum-free medium, ∼ 2-fold higher GREM1 uptake was detected (Suppl. Fig 5). A reduction in GREM1 endocytosis in the presence of BMP2, with more marked GREM1 binding to cell membranes and cell-cell junctions was also detected (Suppl. Fig. 5). These data suggest that protein (or other) components in FBS may be competing with GREM1, thereby reducing cell binding. Conversely, GREM1-BMP2 complexes appear to bind to, but not enter cells in the absence of FBS, suggesting that unknown ingredients of FBS are required for full uptake of GREM1- BMP2 complexes.

The absolute requirement for BMP2 binding to GREM1 for cellular uptake was demonstrated by the dramatic reduction in the uptake of the non-BMP2 binding GREM1^Mut^ protein even in the absence of exogenous BMP2 (Fig. 9. 10). This attenuation of uptake was not due to the stability of the GREM1^Mut^ protein, as equal levels of GREM1^Mut^ were detected in the CM pre- and post-incubation (Fig. 9E, F). Several groups have identified receptor tyrosine kinases such as VEGFR2, EGFR and FGFR1 as targets for GREM1 (Metola et al., 2010), (Park et al., 2020), (Cheng et al., 2022). To date, it has not been clear whether the ability of GREM1 to bind to BMP ligands modulates this RTK signaling. The GREM1^Mut^, which we clearly demonstrated does not bind to BMP2 in both HCT116 (Fig. 9C) and C2C12 cells (Suppl. Fig. 1C) may be a valuable reagent to further refine our understanding of GREM1-RTK signaling in cancer and other diseases.

## Supplemental Video file

HCT116 cells were treated with GREM1-FITC (1 μg/ml) overnight. Fluorescence images were captured every 30 min on a Nikon 6D live-cell imaging microscope. Exported images at the indicated times were used to compile a video to demonstrate GREM1-FITC uptake and localization over time. Scale bars, 50 μm.

## Disclosure Statement

Apart from Fig. 9B, no generational AI software was used to generate figures or text for this manuscript.

## Data Availability

All data used in this manuscript will be made available via a publicly available database, unless protected by licence or other legal barriers.

## Acknowledgements

The authors thank Dr. Ileana Micu and Dr. Ryan Delaney in the imaging core technology unit for assistance with microscopy, Dr. Emma Evergren and Dr. Gunnar Schroeder for advice on organelle staining, Dr. Yue Su and Dr. Anna Krasnodembskaya for assistance with EV isolation, and other members of the technical and academic team at WWIEM for advice and support.

---

Supplemental Figure 1. B**M**P2 **and BMP4 activity is inhibited by GREM1-FITC and GREM1^WT^-mCherry, but not GREM1^MUT^-mCherry.** A. HeLa cells were cultured in medium containing 1 % FBS overnight, followed by 3 h serum-free treatment. Cells were then treated for 60 min at 37 °C with either vehicle (4 mM HCl and 0.1 % BSA) or 5 ng/mL rhBMP2 in the presence of either 25 ng/mL unconjugated rhGREM1, unconjugated FITC, or FITC-GREM1. Pre-incubation of FITC-GREM1 with vehicle or 5 ng/mL rhBMP2 was performed for 15 min at 37 °C. Protein was then extracted and analysed by Western blotting using pSMAD1/5/8 and total SMAD1 antibodies. β-actin was used as loading control. Figure is representative of n=3 independent experiments. B. HEK293 cells were transfected with plasmids expressing mCherry, GREM1^WT^-mCherry and GREM1^MUT^-mCherry. After 24 h, cells were treated with either vehicle (4 mM HCl and 0.1 % BSA) or 5 ng/mL BMP4 for 1 h. Cell lysates were analyzed by Western blot using antibodies reactive to pSMAD1/5/8, total SMAD1, hGREM1 and mCherry. β-actin was used as a loading control. Data representative of n=3 independent experiments carried out in duplicate. C. C2C12 cells stably transfected with BMP response elements (C2C12-BREs, Inman ref) treated with increasing amounts of CM containing GREM1^WT^-mCherry or GREM1^MUT^-mCherry overnight (16 h) prior to lysis. Luciferase activity was assessed in 96-well plate format. Icons are representative of the average of duplicate parallel tests, carried out in triplicate wells. Luciferase response curves were generated using GraphPad Prism to determine half-maximal inhibitory values (IC_50_). Data are plotted as mean ± SEM. One-way ANOVA followed by Bonferroni post-hoc test was used for statistical analysis. **, p<0.01.

Supplemental Figure 2. G**R**EM1**-FITC internalization in HCT116 cells.** HCT116 cells were treated overnight with 1 μg/mL GREM1-FITC. Bright field and fluorescence “snapshot” images were captured every 30 min using a Nikon 6D live cell imaging microscope, and a live cell imaging video was compiled. Exported images at the indicated time-points demonstrate GREM1-FITC localization. The zoomed-in sections highlight the detailed localization of GREM1-FITC within the cells. Scale bar, 50 μm.

Supplemental Figure 3. G**R**EM1 **does not localize to mitochondria, endoplasmic reticulum or Golgi apparatus.** HeLa cells were seeded to a sparse density on Ibidi µ-Slides. After adherence, cells were treated with 1 μg/mL GREM1-FITC (green) overnight in complete medium before being stained with (a) 200 nM Mitotracker (red) to visualize mitochondria, (b) anti-Calreticulin (red) to visualize endoplasmic reticulum or (c) anti-GM130 (red) to visualize Golgi apparatus in serum-free growth medium for 30 min at 37 °C. Cells were then stained with DAPI (blue) to visualize the nucleus. Slides were then images at 40 x magnification on the confocal microscope Leica SP5 as described in Methods. Scale bars represent 50 μm (a), 25 μm (b, c) or 25 μm (a), 10 μm (b, c) in the magnified panels. Data representative of n=3 independent experiments.

Supplemental Figure 4. G**R**EM1 **partially localizes to lysosomes but not recycling endosomes.** HeLa cells were treated with 0.5 μg/mL GREM1-FITC overnight in complete medium. Cells were then washed with PBS and fixed with 4 % PFA (w/v) before staining with (A) anti-LAMP-1 to visualize lysosomes (red) or (B) anti-RAB11 to stain for recycling endosomes, and DAPI (blue) to visualize nuclei. Slides were then imaged at 100 x magnification on a Leica SP5 confocal microscope. Scale bars represent 25 μm or 10 μm in the magnified images. Data representative of n=3 independent experiments.

Supplemental Figure 5. B**M**P2**-stimulated GREM1 uptake requires FBS.** Conditioned medium (CM) from HEK293 cells transfected with GREM1^WT^-mCherry was added to HCT116 cells in the presence of vehicle (4 mM HCl and 0.1 % BSA) or 200 ng/mL BMP2 overnight (16 h) in either serum-free (-FBS) or 10 % FBS (+FBS)-containing medium. Cells were fixed and stained with DAPI before imaging using a Leica DMi8 microscope at 20 x magnification.

## Notes

### Competing Interest Statement

The authors have declared no competing interest.

