## Supplementary material for "Identification of a novel GREMLIN1 uptake pathway in epithelial cells that requires BMP binding": Figures

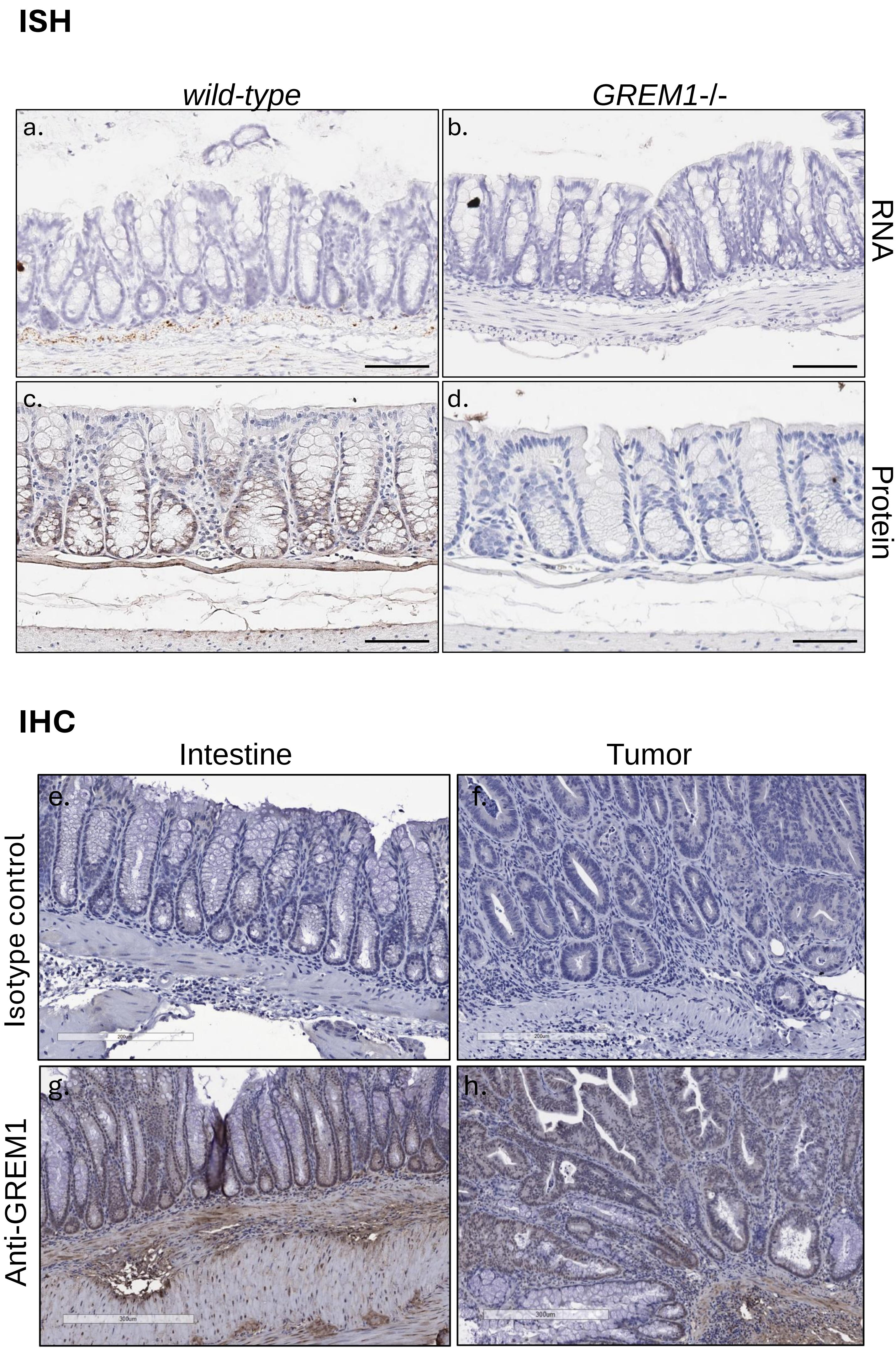
ISH
wild-type
GREM1-/-
a.
b.
RNA
c.
d.
Protein
IHC
Intestine
Tumor
e.
f.
Isotype control
g.
h.
Anti-GREM1

Figure 1.

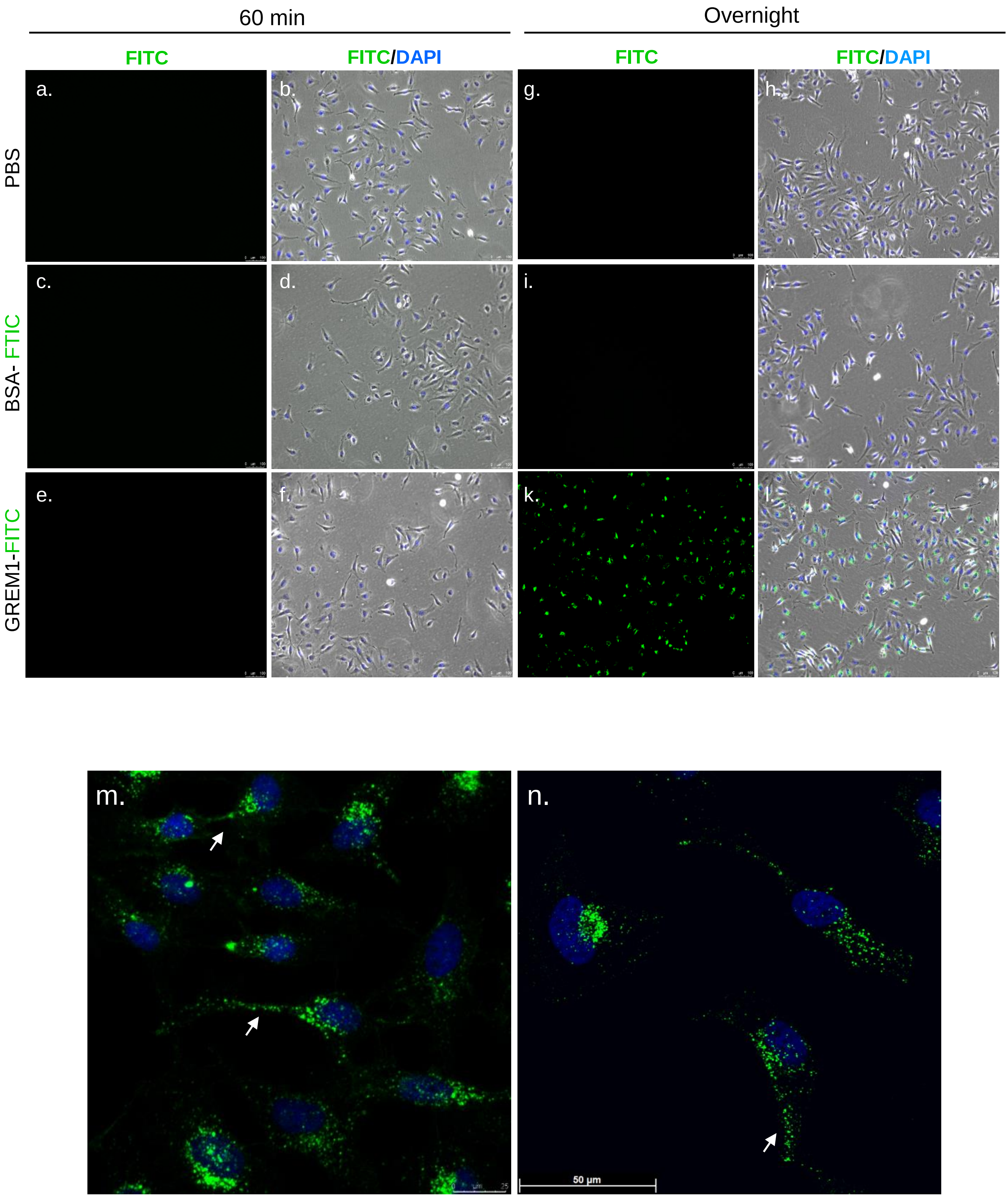
60 min
Overnight
FITC
FITC/DAPI
FITC
FITC/DAPI
PBS
BSA- FTIC
GREM1-FITC
a.
b.
c.
d.
e.
f.
g.
h.
i.
j.
k.
l.
m.
n.
50 µm

Figure 2.

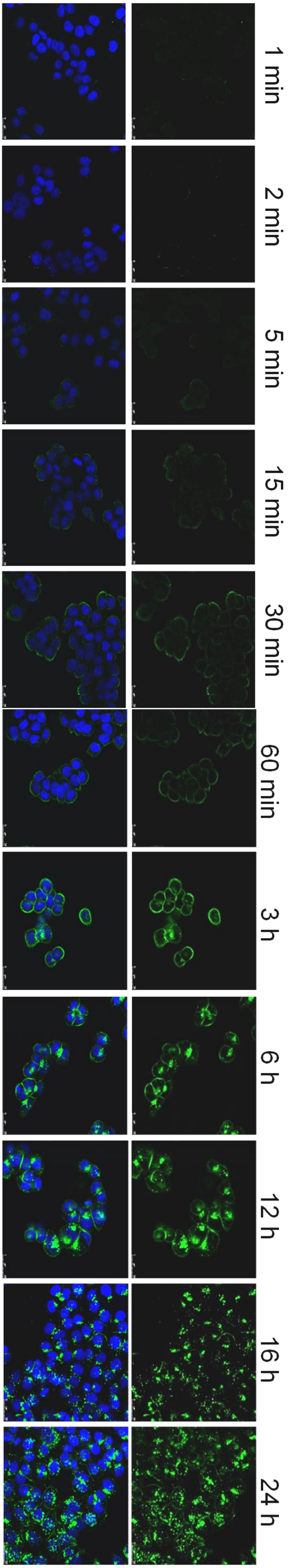


Figure 3.

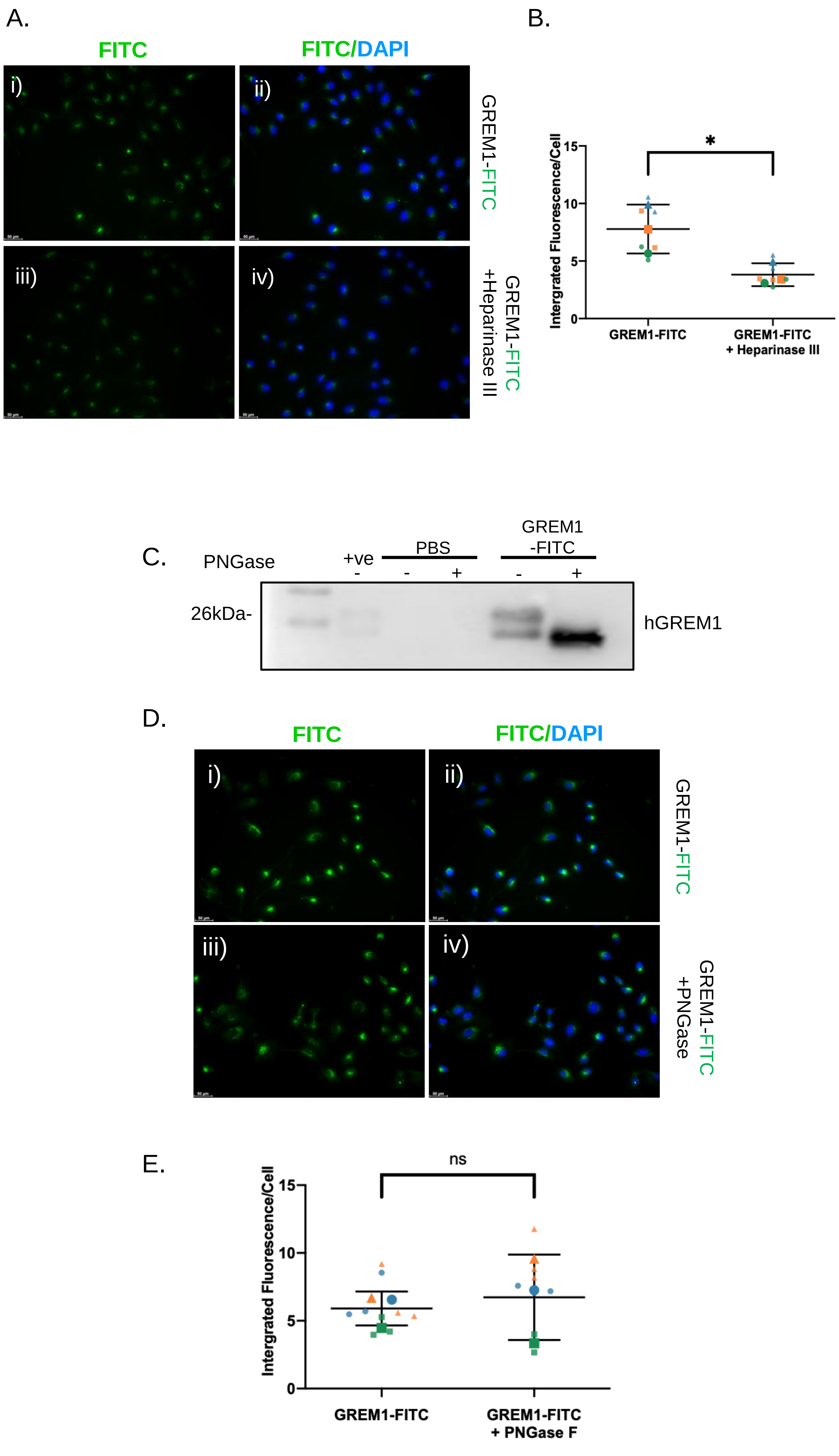

A.
FITC
FITC/DAPI
i)
ii)
iii)
iv)
GREM1-FITC
GREM1-FITC +Heparinase III
B.
Intergrated Fluorescence/Cell
15
10
5
0
*
GREM1-FITC
GREM1-FITC + Heparinase III
C.
PNGase
+ve
PBS
GREM1 -FITC
-
-
+
-
+
26kDa-
hGREM1
D.
FITC
FITC/DAPI
i)
ii)
iii)
iv)
GREM1-FITC
GREM1-FITC +PNGase
E.
Intergrated Fluorescence/Cell
15
10
5
0
ns
GREM1-FITC
GREM1-FITC + PNGase F


Figure 4.

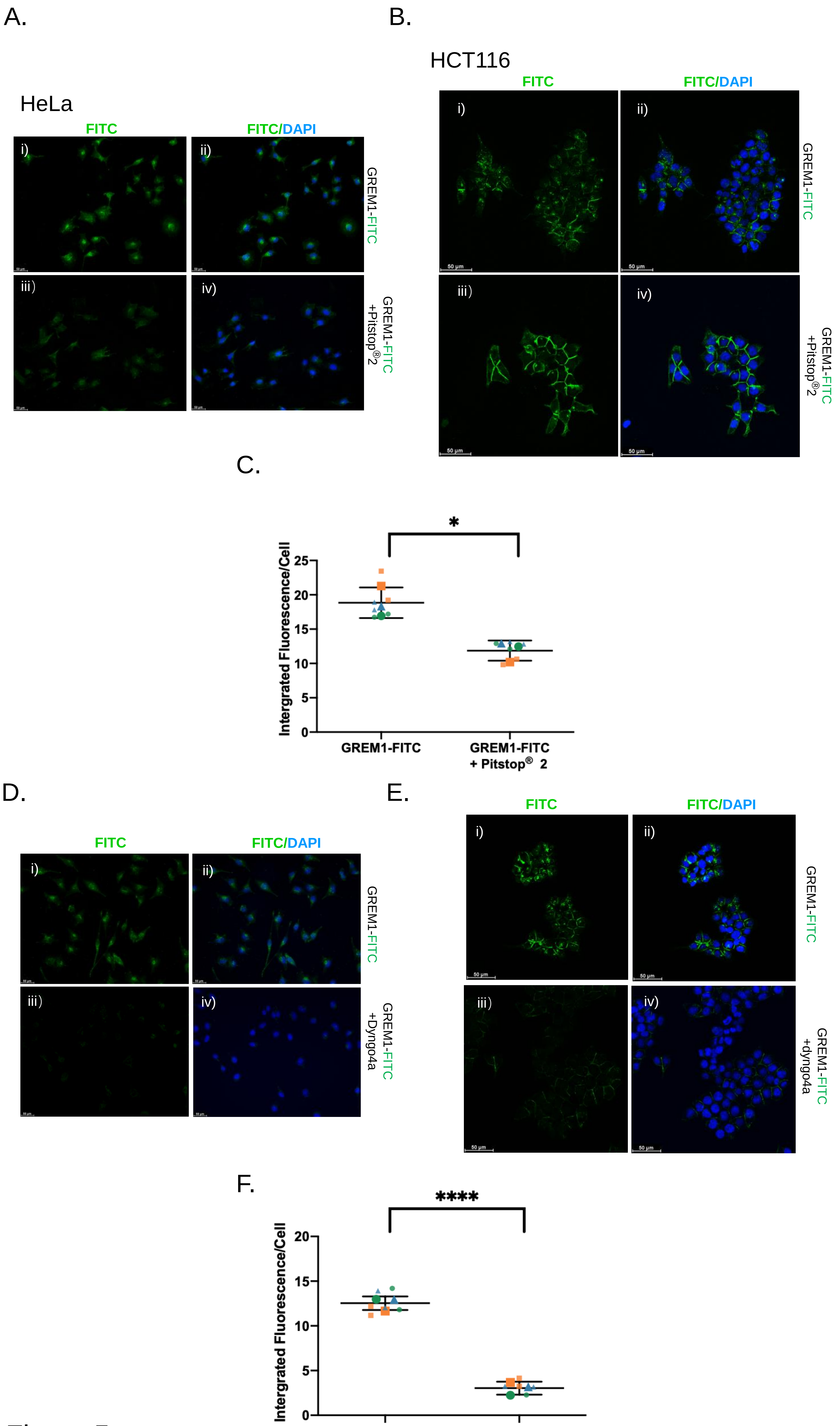
A.
HeLa
FITC
FITC/DAPI
i)
ii)
iii)
iv)
GREM1-FITC
GREM1-FITC +Pitstop® 2
B.
HCT116
FITC
FITC/DAPI
i)
ii)
iii)
iv)
GREM1-FITC
GREM1-FITC +Pitstop® 2
50 µm
C.
*
Intergrated Fluorescence/Cell
25
20
15
10
5
0
GREM1-FITC
GREM1-FITC + Pitstop® 2
D.
FITC
FITC/DAPI
i)
ii)
iii)
iv)
GREM1-FITC
GREM1-FITC +Dyngo4a
E.
FITC
FITC/DAPI
i)
ii)
iii)
iv)
GREM1-FITC
GREM1-FITC +dyngo4a
50 µm
F.
****
Intergrated Fluorescence/Cell
20
15
10
5
0
GREM1-FITC
GREM1-FITC + Dyngo 4a

Figure 5.

A.

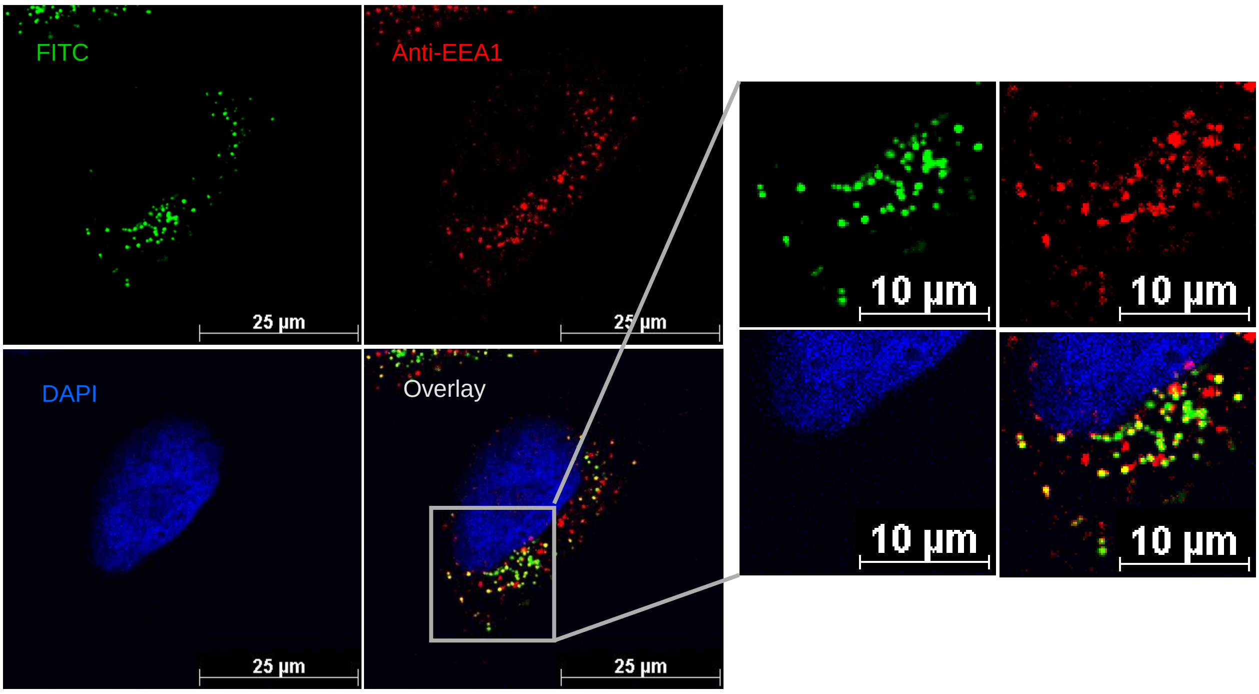
FITC
Anti-EEA1
DAPI
Overlay
25 μm
25 μm
25 μm
25 μm
10 μm
10 μm
10 μm
10 μm

B.

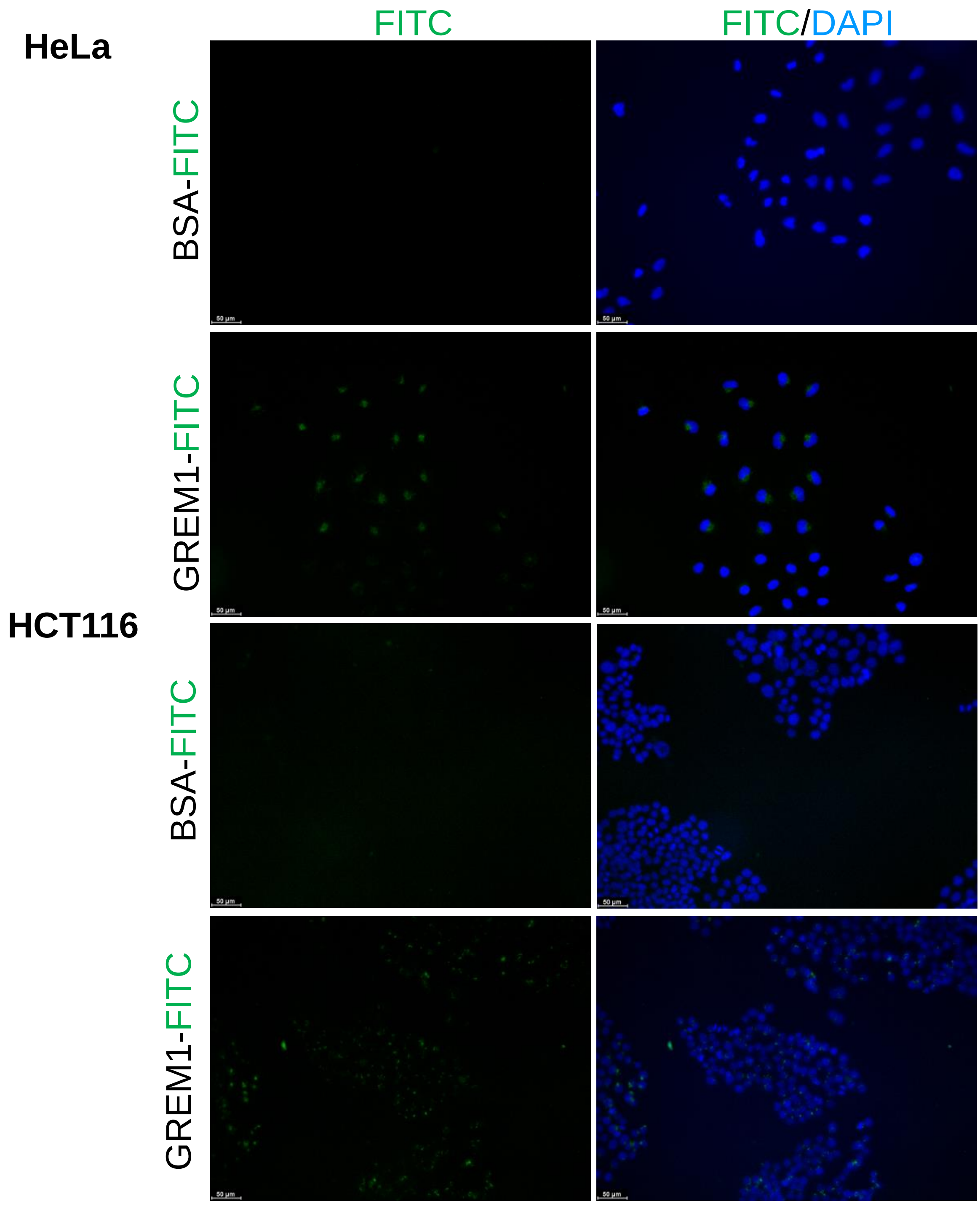
FITC
FITC/DAPI
HeLa
BSA-FITC
GREM1-FITC
HCT116
BSA-FITC
GREM1-FITC

Figure 6.

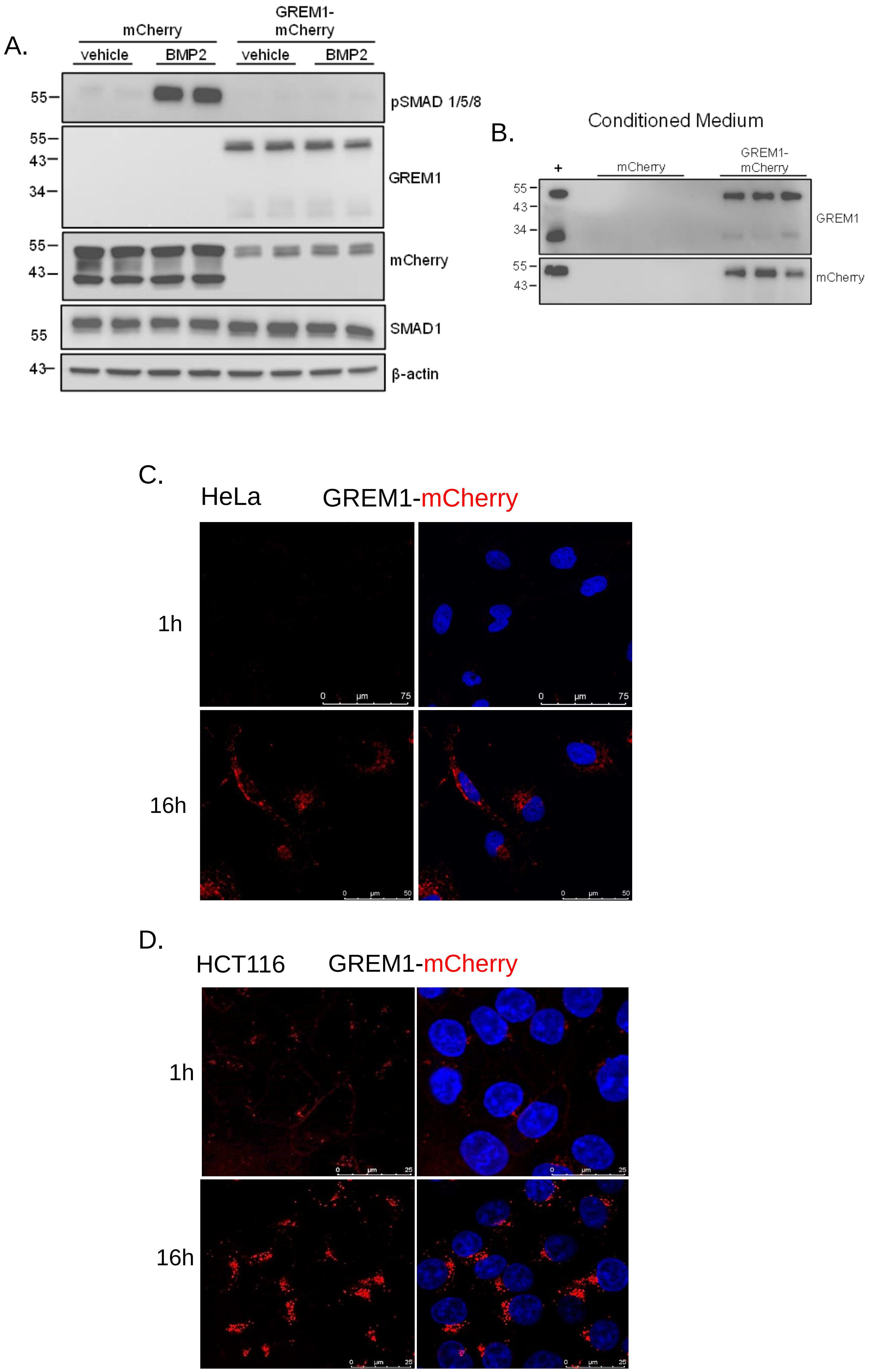
A.
mCherry
GREM1-
mCherry
vehicle
BMP2
vehicle
BMP2
55
43
34
pSMAD 1/5/8
GREM1
mCherry
SMAD1
β-actin
B.
Conditioned Medium
+
mCherry
GREM1-
mCherry
GREM1
mCherry
C.
HeLa
GREM1-mCherry
1h
16h
D.
HCT116
GREM1-mCherry
1h
16h

Figure 7.

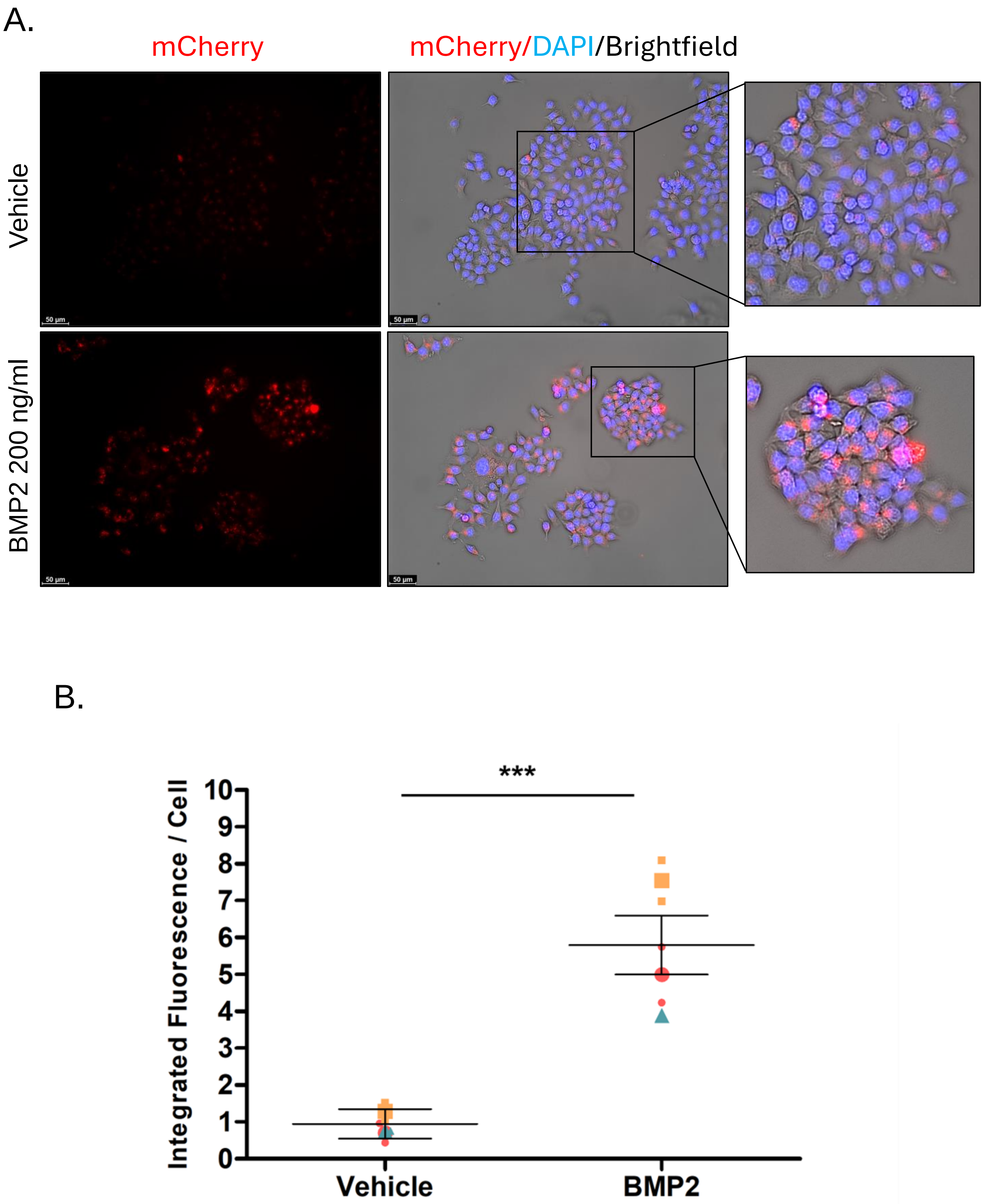
A.
mCherry
mCherry/DAPI/Brightfield
Vehicle
BMP2 200 ng/ml
B.
***
Integrated Fluorescence / Cell
10
9
8
7
6
5
4
3
2
1
0
Vehicle
BMP2

Figure 8.

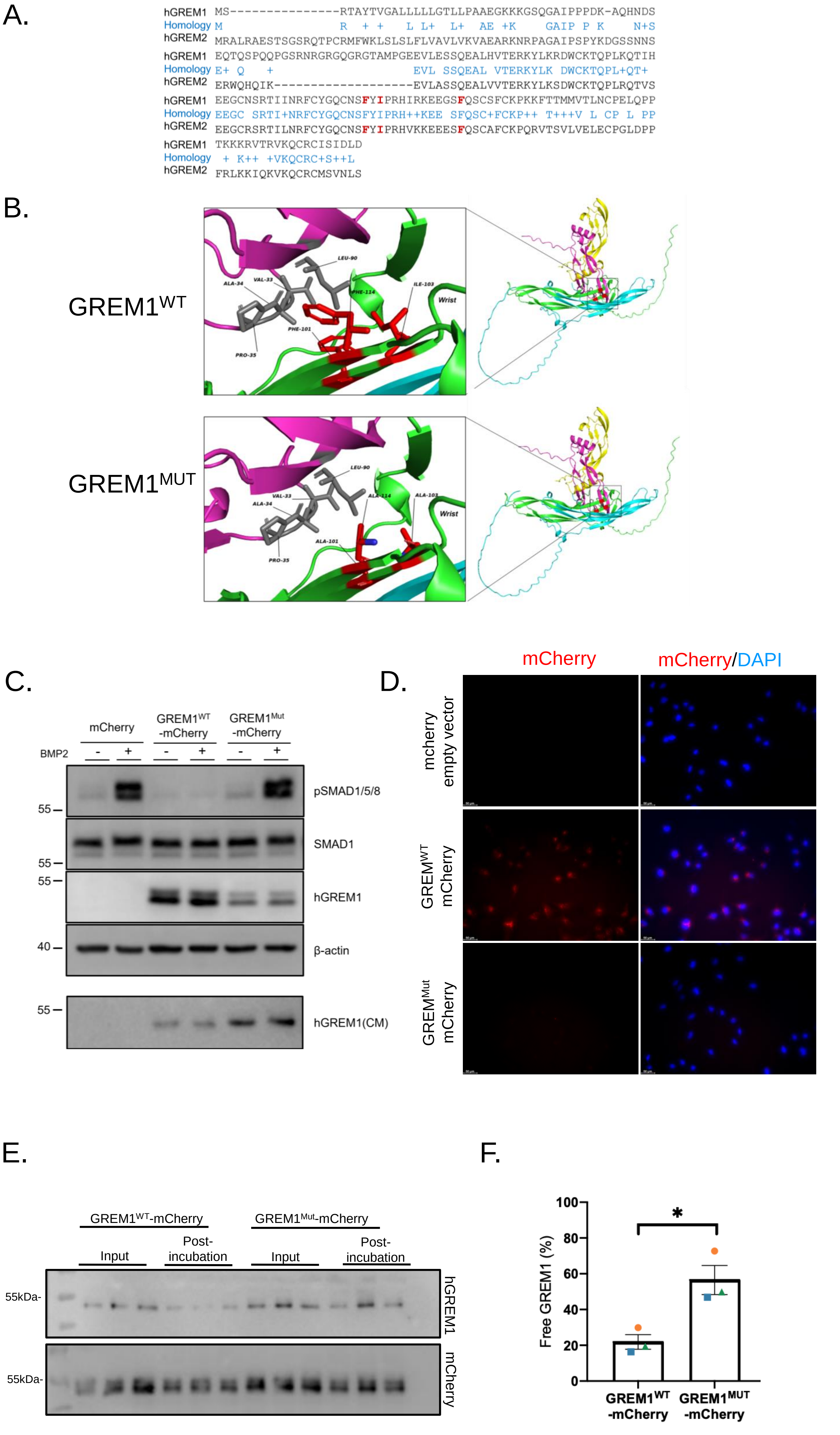
A.
hGREM1 MS---------------RTAYTVGALLLLLGTLLPAAEGKKKGSQGAIPPPDK-AQHNDS
Homology M R + + L L+ L+ AE +K GAIP P K N+S
hGREM2 MRALRAESTSGSRQTPCRMFWKLSLSLFLVAVLVKVAEARKNRPAGAIPSPYKDGSSNNS
hGREM1 EQTQSPQQPGSRNRGRGQGRGTAMPGEEVLESSQEALHVTERKYLKRDWCKTQPLKQTIH
Homology E+ Q + EVL SSQEAL VTERKYLK DWCKTQPL+QT+
hGREM2 ERWQHQIK------------------EVLASSQEALVVTERKYLKSDWCKTQPLRQTVS
hGREM1 EEGCNSRTIINRFCYGQCNSFYIPRHIRKEEGSFQSCSFCKPKKFTTMMVTLNCPELQPP
Homology EEGC SRTI+NRFCYGQCNSFYIPRH++KEE SFQSC+FCKP++ T+++V L CP L PP
hGREM2 EEGCRSRTILNRFCYGQCNSFYIPRHVKKEEESFQSCAFCKPQRVTSVLVELECPGLDPP
hGREM1 TKKKRVTRVKQCRCISIDLD
Homology + K++ +VKQCRC+S++L
hGREM2 FRLKKIQKVKQCRCMSVNLS
B.
GREM1WT
GREM1MUT
LEU-90
VAL-33
ALA-34
PHE-114
ILE-103
PHE-101
PRO-35
Wrist
ALA-114
ALA-103
ALA-101
C.
mCherry
GREM1WT -mCherry
GREM1Mut -mCherry
BMP2
- + - + - +
55
40
pSMAD1/5/8
SMAD1
hGREM1
β-actin
hGREM1(CM)
D.
mCherry
mCherry/DAPI
mcherry empty vector
GREMWT mCherry
GREMMut mCherry
E.
GREM1WT-mCherry
GREM1Mut-mCherry
Input
Post-incubation
55kDa-
hGREM1
mCherry
F.
Free GREM1 (%)
0
20
40
60
80
100
*
GREM1WT -mCherry
GREM1MUT -mCherry

Figure 9.

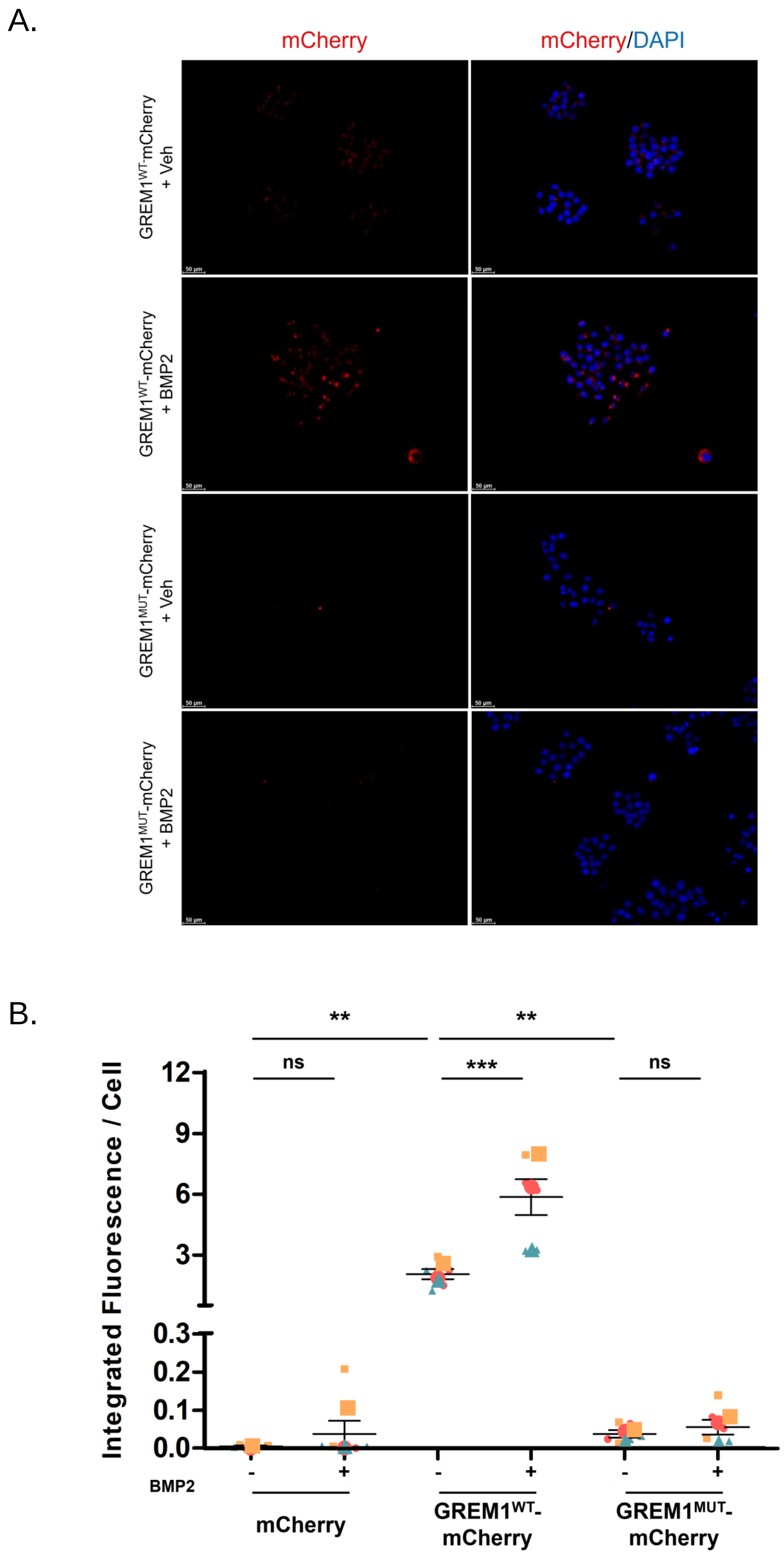

A.
mCherry
mCherry/DAPI
GREM1WT-mCherry + Veh
GREM1WT-mCherry + BMP2
GREM1MUT-mCherry + Veh
GREM1MUT-mCherry + BMP2
B.
**
**
ns
***
ns
Integrated Fluorescence / Cell
12
9
6
3
0.3
0.2
0.1
0.0
BMP2
-
+
-
+
-
+
mCherry
GREM1WT-mCherry
GREM1MUT-mCherry


Figure 10.

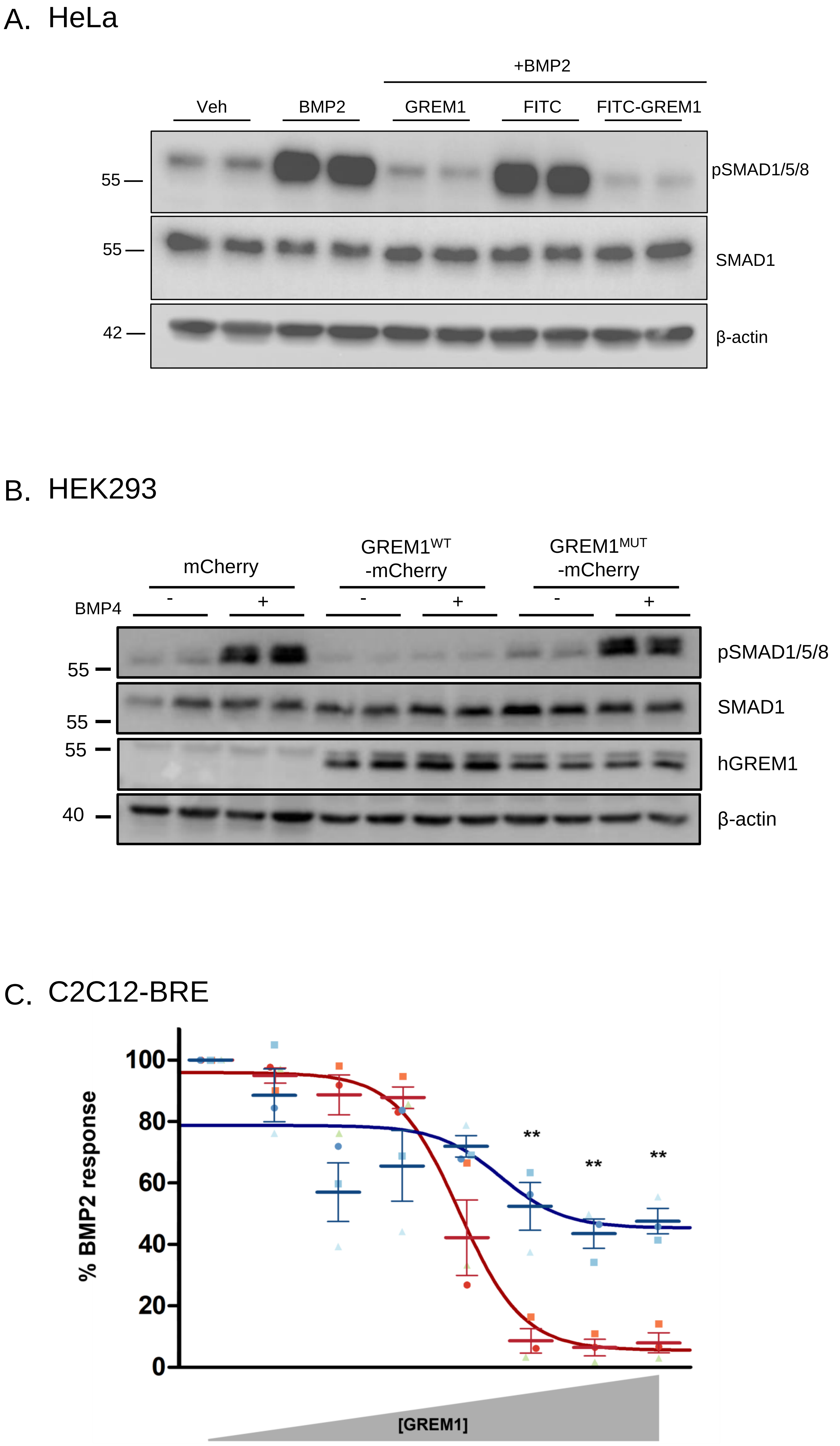


Supplemental Figure 1.

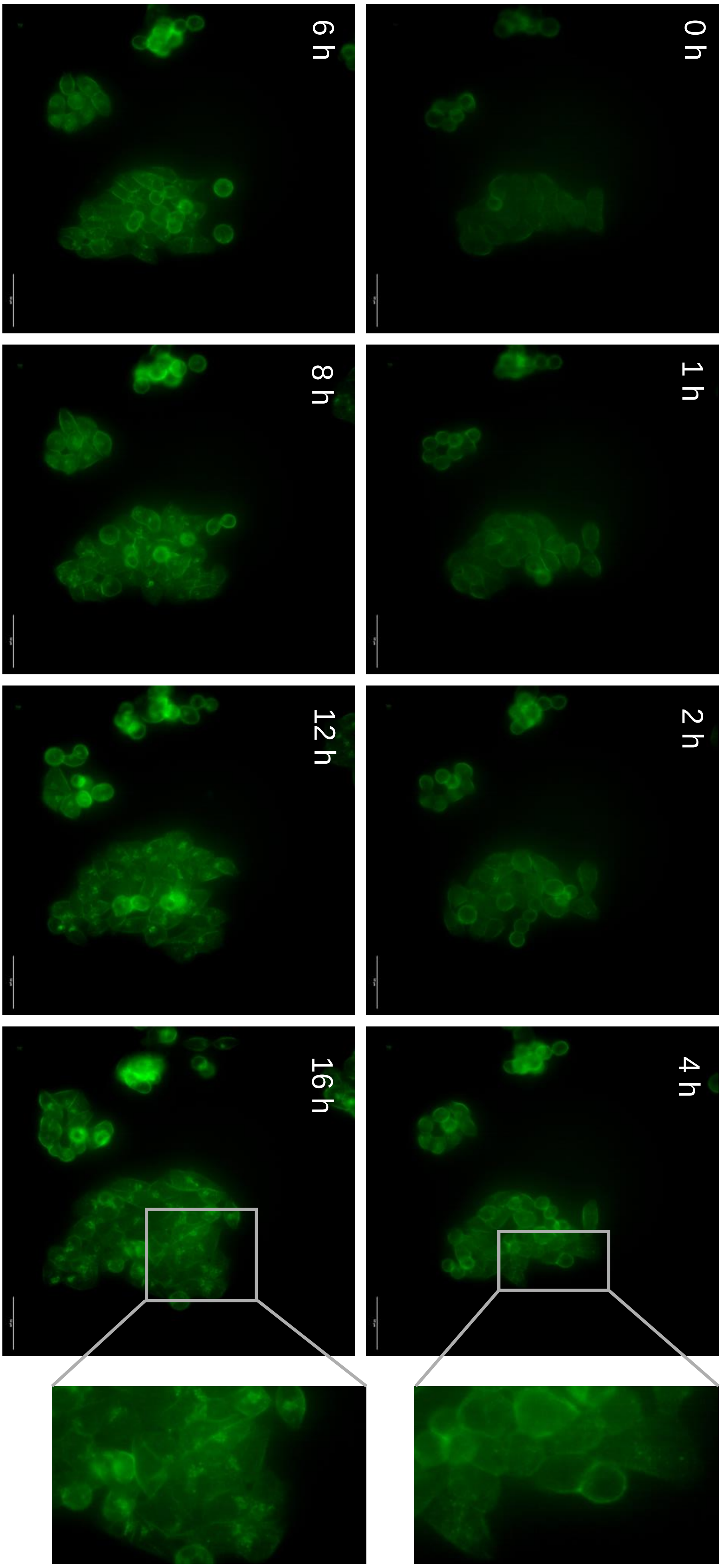


Supplemental Figure 2.

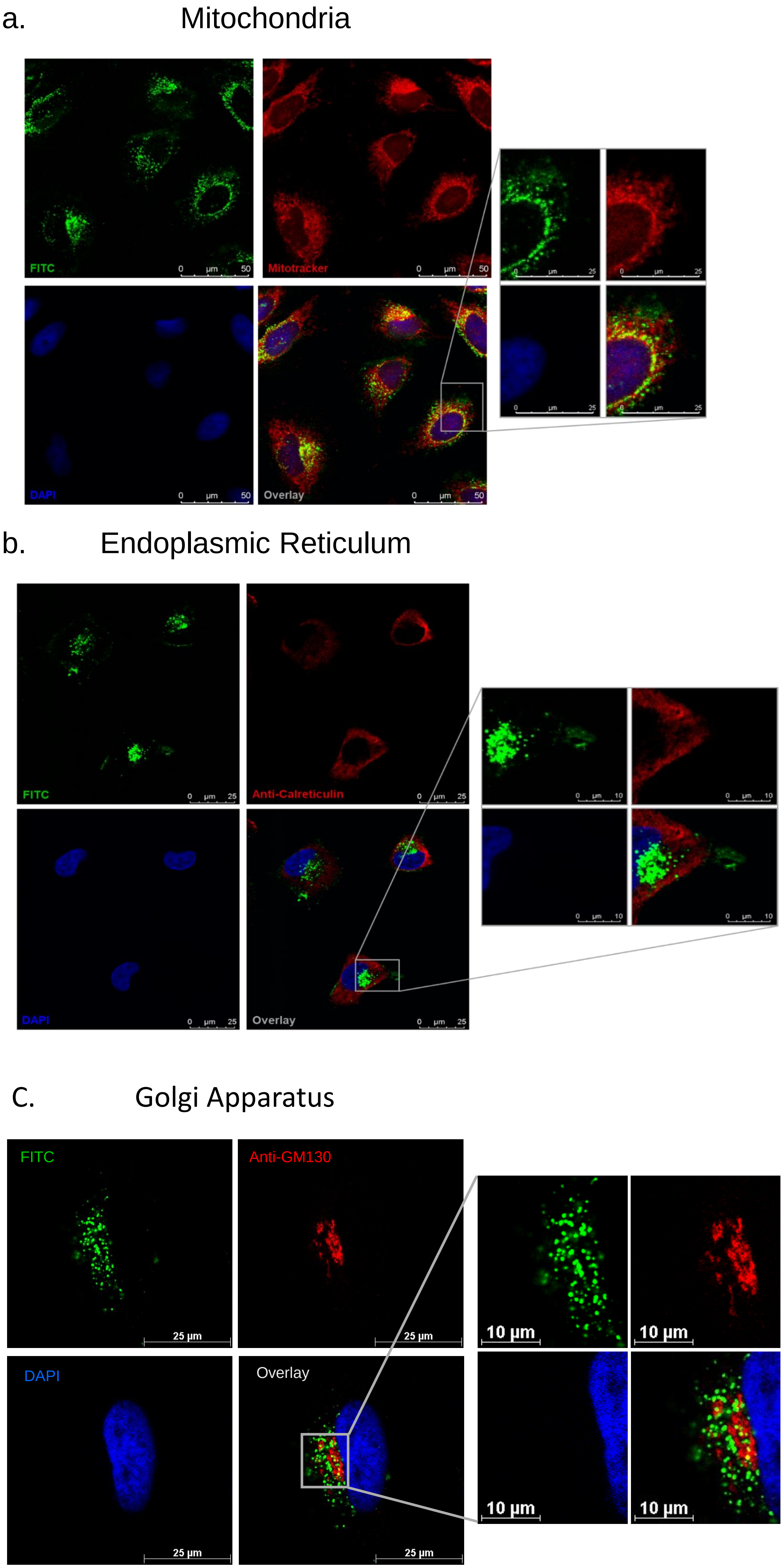


Supplemental Figure 3.

A. Lysosomes

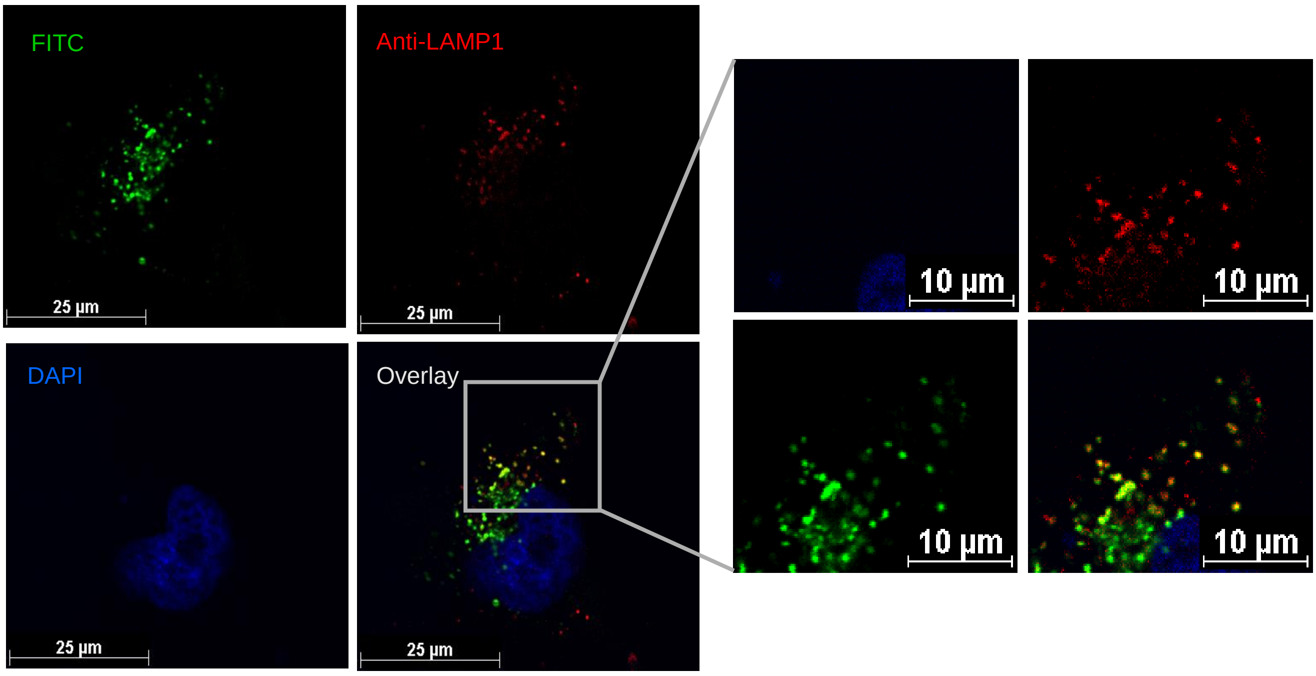


B. Recycling endosomes

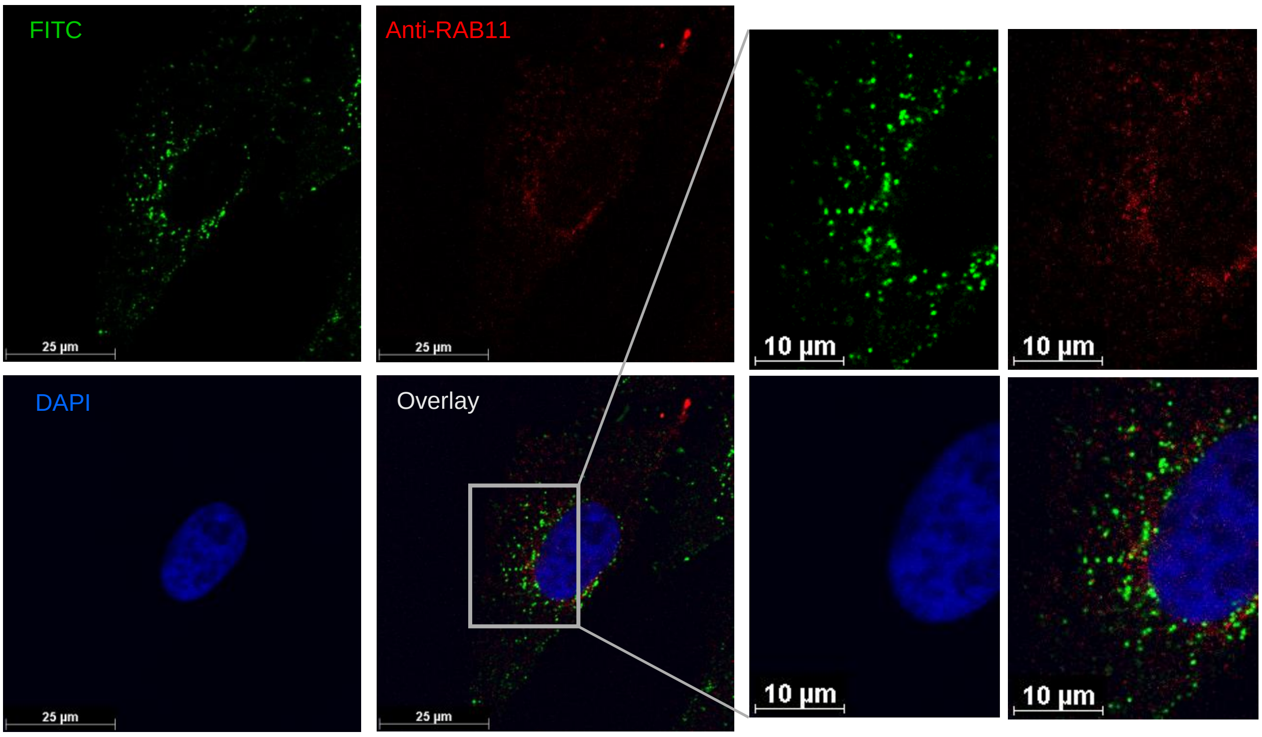


Supplemental Figure 4.

A.

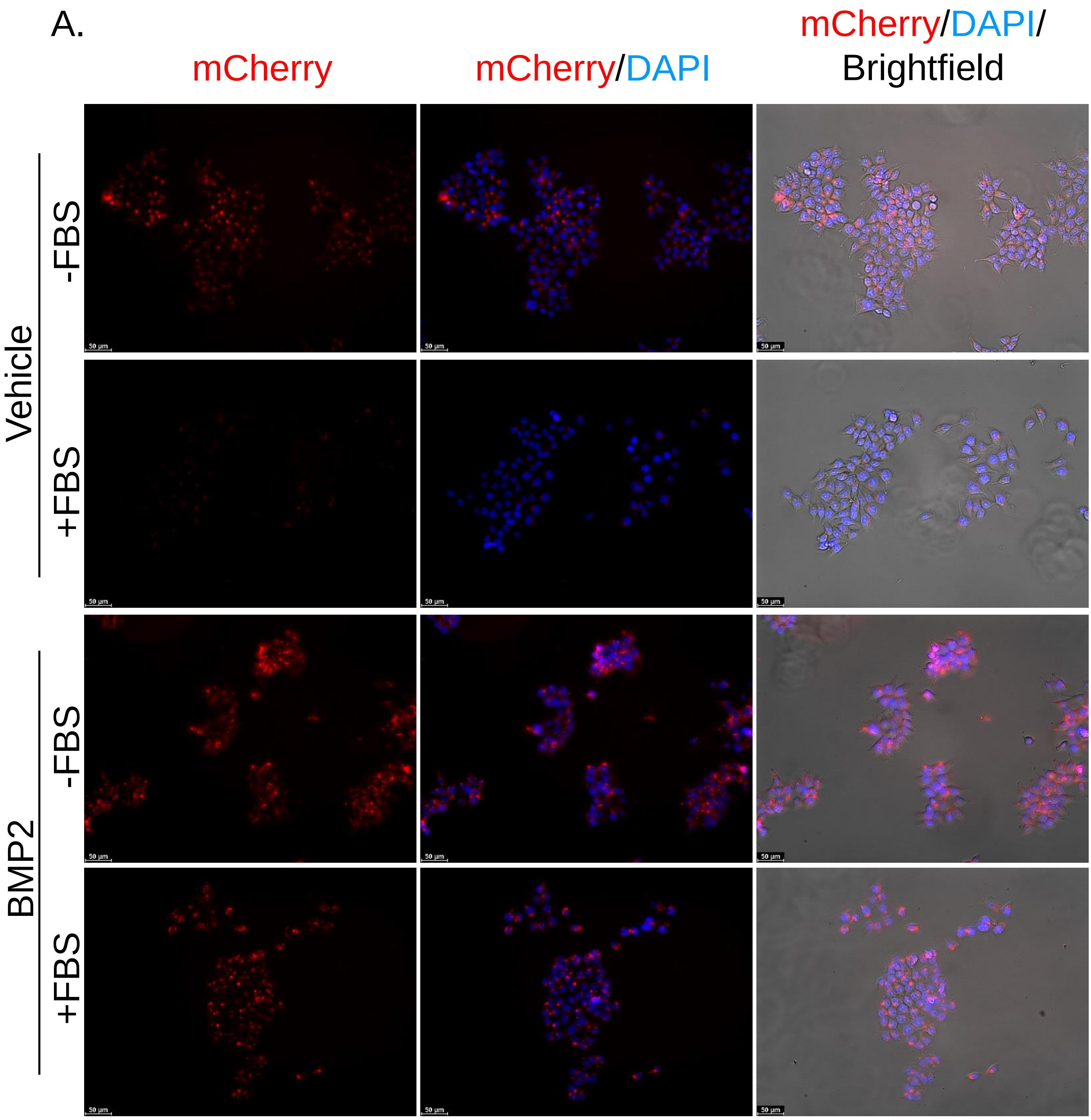


B.

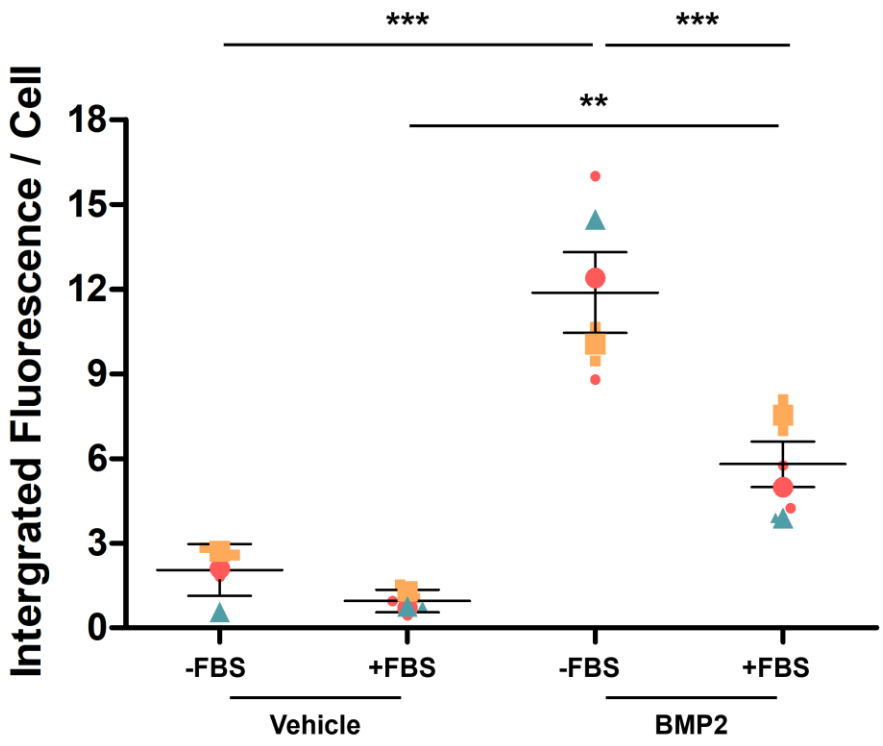


Supplemental Figure 5.

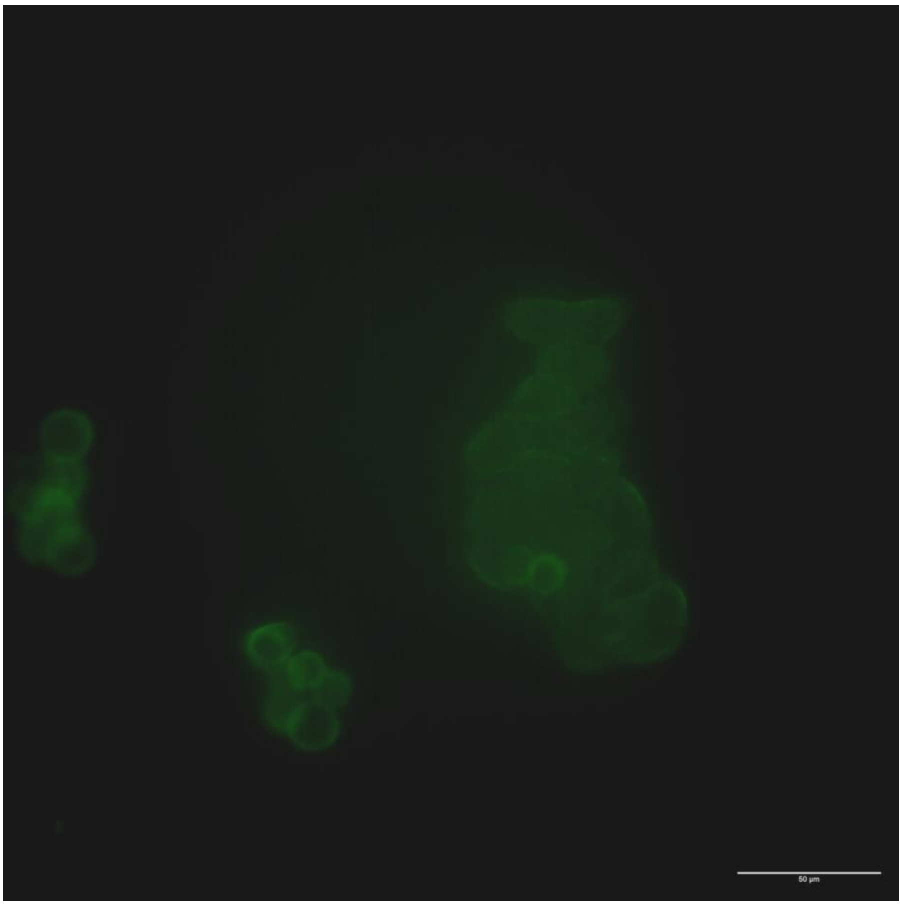


Supplemental Video File
